# Helical metaphase chromatid coiling is conserved

**DOI:** 10.1101/2021.09.16.460607

**Authors:** Ivona Kubalová, Amanda Souza Câmara, Petr Cápal, Tomáš Beseda, Jean-Marie Rouillard, Gina Marie Krause, Helena Toegelová, Axel Himmelbach, Nils Stein, Andreas Houben, Jaroslav Doležel, Martin Mascher, Hana Šimková, Veit Schubert

**Affiliations:** Leibniz Institute of Plant Genetics and Crop Plant Research (IPK) Gatersleben, D-06466 Seeland, Germany; Institute of Experimental Botany of the Czech Academy of Sciences, Centre of the Region Haná for Biotechnological and Agricultural Research, Šlechtitelů 31, 77900 Olomouc, Czech Republic; Daicel Arbor Biosciences, Ann Arbor, MI, USA; Chemical Engineering Department, University of Michigan, Ann Arbor, MI, USA; Center of Integrated Breeding Research (CiBreed), Department of Crop Sciences, Georg-August-University, Von Siebold Str. 8, 37075 Göttingen, Germany; German Centre for Integrative Biodiversity Research (iDiv) Halle-Jena-Leipzig, Leipzig, Germany

**Keywords:** chromatin condensation, chromonema, chromosome conformation capture sequencing (Hi-C), flow cytometric sorting, helical chromatid structure, *Hordeum vulgare*, mitotic metaphase chromosome, oligo-fluorescence in situ hybridization (oligo-FISH), polymer simulation, sister chromatid exchange (SCE), structured illumination microscopy (SIM)

## Abstract

The higher-order organization of metaphase chromosomes has been debated for almost 140 years. Classical light and electron microscopy studies suggested that chromatids are composed of helically organized chromatin fibers (chromonemata). Non-helical models were also recently proposed. We studied chromosome organization in barley using cutting-edge approaches and obtained evidence for a helically arranged 400-nm chromatin fiber representing the chromonema within chromatid arms. The number of turns is positively correlated with arm length. Turn size and chromatin density decrease towards the telomeres. Due to their specialized functions, the helical organization of centromeres and nucleolus-organizing regions is interrupted by several thinner, straight chromatin fibers. A comparison with previously published data indicates that the helical turning of metaphase chromatid arms is a conserved feature of large eukaryotic chromosomes.

## Introduction

Cell growth, division, and differentiation are accompanied by drastic changes in chromatin structure to facilitate DNA replication, repair, transcription, and segregation. The most striking changes are observed during mitosis and meiosis, when the chromatin of interphase chromosome domains becomes compacted ∼1000-fold to form rod-like structures. After replication, during the S phase of the cell cycle, the chromosomes consist of a pair of genetically identical chromatids (Paulson and Laemmli 1977; Marsden and Laemmli 1979; Earnshaw and Laemmli 1983; Maeshima et al. 2005). Individual chromosomes can most easily be observed during nuclear division. While some species have holocentric chromosomes (containing elongated centromeres), most eukaryotes have monocentric chromosomes characterized by a distinct primary constriction to which spindle fibers attach for proper chromatid segregation during anaphase (Musacchio and Desai, 2017; Schubert et al., 2020). Some chromosomes also harbor a nucleolus organizer region (NOR), a 45S rDNA cluster forming a secondary constriction on metaphase chromosomes (McClintock, 1934; Schöfer and Weipoltshammer, 2018). Long-standing questions about chromatin structure remain, including how DNA is compacted into rod-shaped mitotic and meiotic chromosomes, which principles govern folding, and whether such principles apply to different species.

Chromatin in interphase nuclei comprises 10-nm nucleosome fibers formed by ∼147 bp of DNA wound around an octamer of nucleosome histones. The nucleosomes are connected by linker DNA like “beads-on-a-string” (Thoma et al. 1979). However, little is known about how nucleosome fibers fold into higher-order structures. Using purified 10 nm fibers, Finch and Klug (1976) observed the formation of a 30-nm chromatin fiber, the so-called solenoid. Nevertheless, its existence in native chromatin has not been confirmed, and there is growing evidence that the 30 - nm fiber forms in vitro only in solutions with low salt concentrations. Several studies across various eukaryotes demonstrated that the higher-order structure of chromatin is formed by an irregularly folded 10-nm fiber (Grigoryev et al. 2016; Maeshima et al. 2010a, b, 2016, 2018, 2019; Nishino et al. 2012). The periodical arrangement of chromatin has been questioned as well. Wako et al. (2020) analyzed chromatin fibers with different levels of compactness in human metaphase chromosomes using beam/scanning electron microscopy. Thick (30 nm) fibers were rare, excluding the possibility that they are structural units. In contrast to several existing models (Finch and Klug 1976; Manuelidis and Chen 1990), no periodical structure was found. While most researchers accept the notion that metaphase chromosomes are composed of 10 nm fibers, the higher-order organization remains obscure.

Several models have been proposed to describe the higher-order structure of metaphase chromosomes based on data obtained using a range of molecular and microscopy methods (Beseda et al. 2020). These models are categorized as helical and non-helical. Helical models assume that the chromatin in each sister chromatid at metaphase is arranged in a helix (Manton 1950; Ohnuki, 1965, 1968; Iino 1971; Bak et al. 1977; Sedat and Manuelidis 1978; Marsden and Laemmli 1979; Ris 1981; Rattner and Lin 1985; Nokkala and Nokkala 1986; Boy de la Tour and Laemmli 1988; Saitoh and Laemmli 1994; Sumner 1998; Inaga et al. 2002; Daban 2020; Gibcus et al. 2018; Schloissnig et al. 2021), whereas non-helical models suggest that chromatin is folded within the chromatids without forming a helix (Paulson and Laemmli 1977; Belmont et al. 1987, 1989; Wanner and Formanek 2000; Poirier and Marko 2002; Strukov et al. 2003; Kireeva et al. 2004; Marko 2011; Naumova et al. 2013; Chu et al. 2020).

The helical structure of chromosomes was first observed in the plant *Tradescantia virginica* L. by Baranetzky (1880) and was later supported by Ruch (1949). A similar chromatin organization called chromonema (from chromo- + Greek nēma thread; plural: chromonemata) was discovered in diverse plant species by observing native, physically, and/or chemically treated chromosomes by light microscopy (Nebel 1932a, 1939; Manton 1950; Schvartzman et al. 1978; Nokkala and Nokkala 1985). Interestingly, such chromatin coiling was not found in Drosophila (Strukov et al. 2011), and although Ohnuki et al. (1968) observed helical coiling in mitotic chromosomes of cultured human leukocytes, such a structure was not observed in mammalian cell lines by other groups (Strukov and Belmont 2009; Chu et al. 2020). Furthermore, the helical chromatin arrangement in mitotic chromatids was not observed in three human cell lines using the chromatin conformation capture (3C) methods 5C and Hi-C, which quantify the frequency of genome-wide chromatin contacts (Naumova et al. 2013). On the contrary, Hi-C data obtained from mitotic cells of chicken (Gibcus et al. 2018) and axolotl (Schloissnig et al. 2021) suggested a helical chromosome organization. After reanalyzing mitotic HeLa cell Hi-C data from Naumova et al. (2013) in more detail by deeper sequencing, Gibcus et al. (2018) confirmed the helical structure and suggested that this arrangement might be a common feature among vertebrates.

Advanced microscopy, cytology, and molecular techniques could be used to decipher the higher-order spatial (3D) chromatin organization of mitotic chromosomes at higher resolution. The use of fluorescence dyes to label specific DNA segments and proteins has led to the development of super-resolution microscopy techniques such as structured illumination microscopy (SIM), a sub-diffraction imaging method bridging the resolution gap between light and electron microscopy (Rego et al. 2012). These nanoscopic methods have been successfully used in cell biology (Fornasiero and Opazo 2015), leading to the discovery of structures within plant chromatin (Schubert,2017; Baroux and Schubert 2018; Schubert et al. 2020).

Fluorescence in situ hybridization (FISH) using oligonucleotide probes (oligo-FISH) is an efficient method for examining whole chromosomes and specific regions to investigate karyotype structure, meiotic chromosome behavior, and chromosome evolution via comparative cytogenetic mapping (Aurich-Costa et al. 2007; Han et al. 2015; Braz et al. 2018; Šimoníková et al. 2019; Xin et al. 2020; Li et al. 2021; Hoang et al. 2021). The application of super-resolution microscopy techniques in combination with oligo-FISH led to the characterization of sub-chromosomal structures (Szabo et al. 2018). The arrangement of chromatin in sister chromatids in spontaneously occurring sister chromatid exchanges (SCEs) can be studied via the differential incorporation of DNA base analogs, such as 5-bromodeoxyuridine (BrdU) (Latt 1973) or 5-ethynyl-2′-deoxyuridine (EdU) (Schubert et al., 2011), into chromosomes during replication (Schvartzman 1987; Wilson and Thompson 2007; Schubert et al. 2016a). This method differentially labels sister chromatids (harlequin staining) and can be combined with FISH (Perry and Wolff 1974; Rachel et al. 1991; Jordan et al. 1999).

3C techniques were recently used to determine the contact probabilities of genomic loci with different distances in a linear sequence (Lieberman-Aiden et al. 2009; van Berkum et al. 2010; Kempfer and Pombo 2020; Oluwadare et al. 2019) and to uncover hierarchical chromatin organization (Oluwadare et al. 2019; Szabo et al., 2019). Using these techniques, together with polymer simulations, Gibcus et al. (2018) postulated a helical organization of late prometaphase chromosomes, comprising 200 nm thick fibers with helical turns covering 12 Mb, in chicken.

To decipher the higher-order structure of plant mitotic chromosomes, we used barley (*Hordeum vulgare* L.; 2n=14; 1C=4.88 Gb, with large chromosomes) (Doležel et al. 2018) as a model. Analysis using a combination of metaphase chromosome-derived Hi-C data, oligo-FISH, SCE detection, super-resolution microscopy, and polymer simulation revealed that sister chromatids are composed of chromatin helices of identical handedness. The helical turns cover 20–38 Mb, creating a ∼400 nm thick fiber. Only centromere and NOR regions are free of the helical higher-order organization. Thus, we experimentally proved the classical model of a helical organization of somatic chromosomes for a plant species with large chromosomes.

## Results

### Hi-C indicates that the mitotic chromatid is organized as a helix with variable turn lengths

To analyze the organization of condensed chromosomes, we purified barley metaphase chromosomes by flow cytometric sorting, sequenced the contact pairs by Hi-C analysis, and aligned the sequences against the barley cv. Morex genome assembly version 2 (Morex v2) (Monat et al. 2019). The Hi-C contact matrix for metaphase chromosomes showed a second diagonal parallel to the main diagonal along the length of the chromosome, which faded in the vicinity of the primary and secondary constrictions and telomeres (Figures 1A, S1). The additional diagonal was unique to metaphase chromosomes and was not observed in interphase chromatin, which only showed a perpendicular diagonal indicative of the conformation described by Rabl in 1885 (Rabl, 1885) (Mascher et al. 2017).

**Figure 1.**
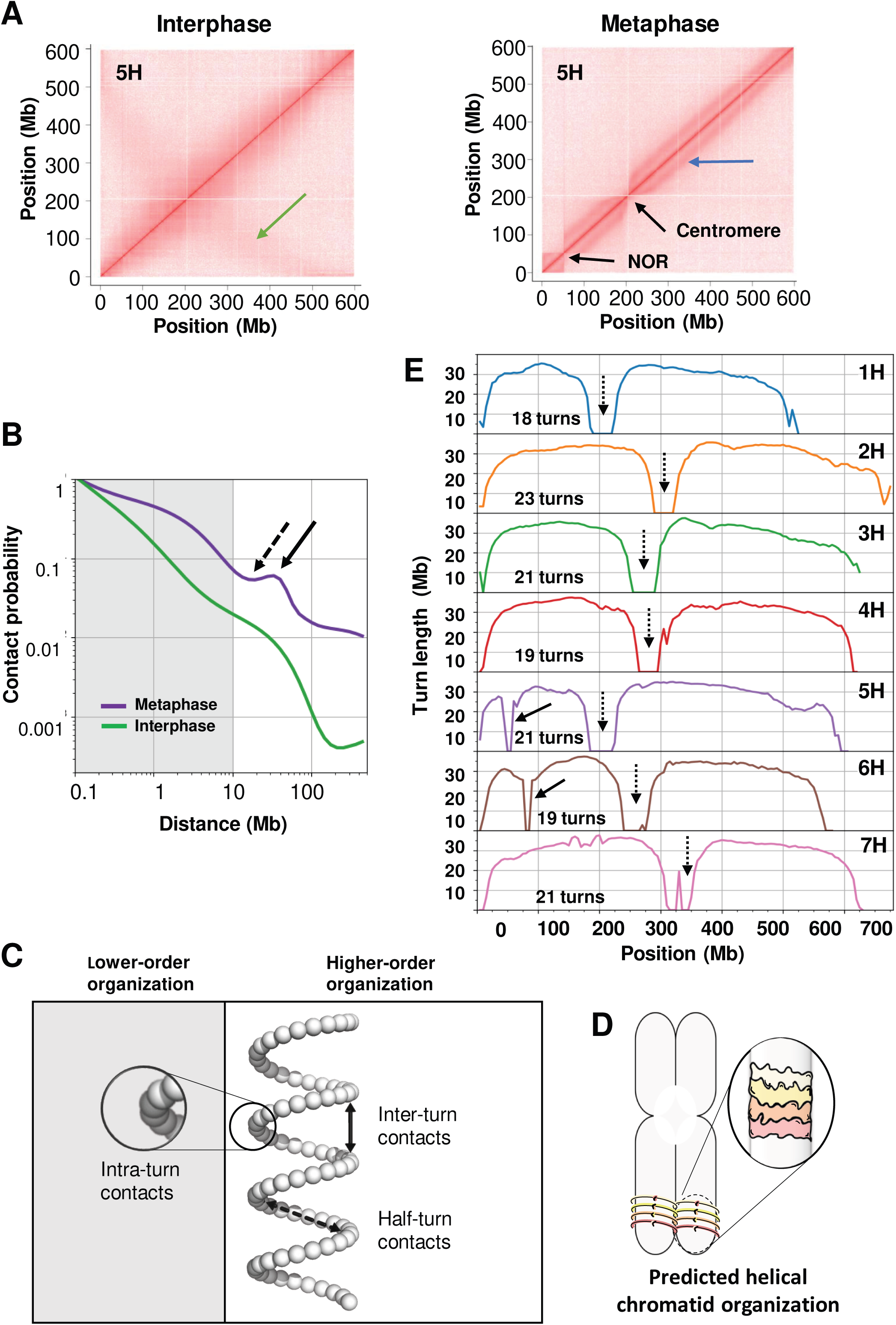
The presence of periodical Hi-C contacts reveals the helical organization of somatic barley metaphase chromosomes. **A)** Hi-C contact matrix of barley chromosome 5H in inter- and metaphase. The anti-parallel diagonal in interphase (green arrow) indicates the Rabl configuration, while metaphase chromosomes show a parallel diagonal (blue arrow), indicating that a periodical contact pattern is missing around the centromere and NOR (black arrows). **B)** Average Hi-C contact probabilities of all chromosomes as a function of the distance between two contacting sites in meta- and interphase. The characteristic bump (solid arrow) relates to the second diagonal observed in the Hi-C metaphase contact matrix (blue arrow in **(A)**. The bump follows a decrease at half of its position (dashed arrow). This pattern indicates an arrangement into turns, where sites distant by one turn (solid arrow) are more likely to come in contact than sites a half-turn apart (dashed arrow). The bump position indicates the turn length. Intra-turn contacts below 10-Mb distance (gray area) indicate the lower-order chromatin organization, where nested loops are packed within each turn (Gibcus et al. 2018). **C)** The three contact types observed with different contact probability patterns **(B)** are represented in a helical arrangement. Sites separated by one turn are closer to each other (solid arrow, inter-turn contacts) than sites separated by a half-turn (dashed arrow, half-turn contacts), and contacts between adjacent loops (inset) constitute a lower-order organization with higher contact probability (gray region). **D)** Schema of a metaphase chromosome arranged into turns, as inferred from the Hi-C matrix **(A)** and the contact probability **(B)**. The color gradient along the turns relates to the linear genomic sequence. **E)** Turn length along all chromosomes based on the local contacts observed in the Hi-C experiments (Table S1). Positions of primary (dashed arrows) and secondary (solid arrows) chromosome constrictions can be recognized by zero turn length. All interstitial regions show similar turn lengths, decreasing towards the telomeres. Abrupt peaks, such as in chromosome 7H at 160, 195, and 330 Mb, relate to gaps or assembly errors in the genome sequence. The total turn number per chromosome is indicated.

**Table 1:**
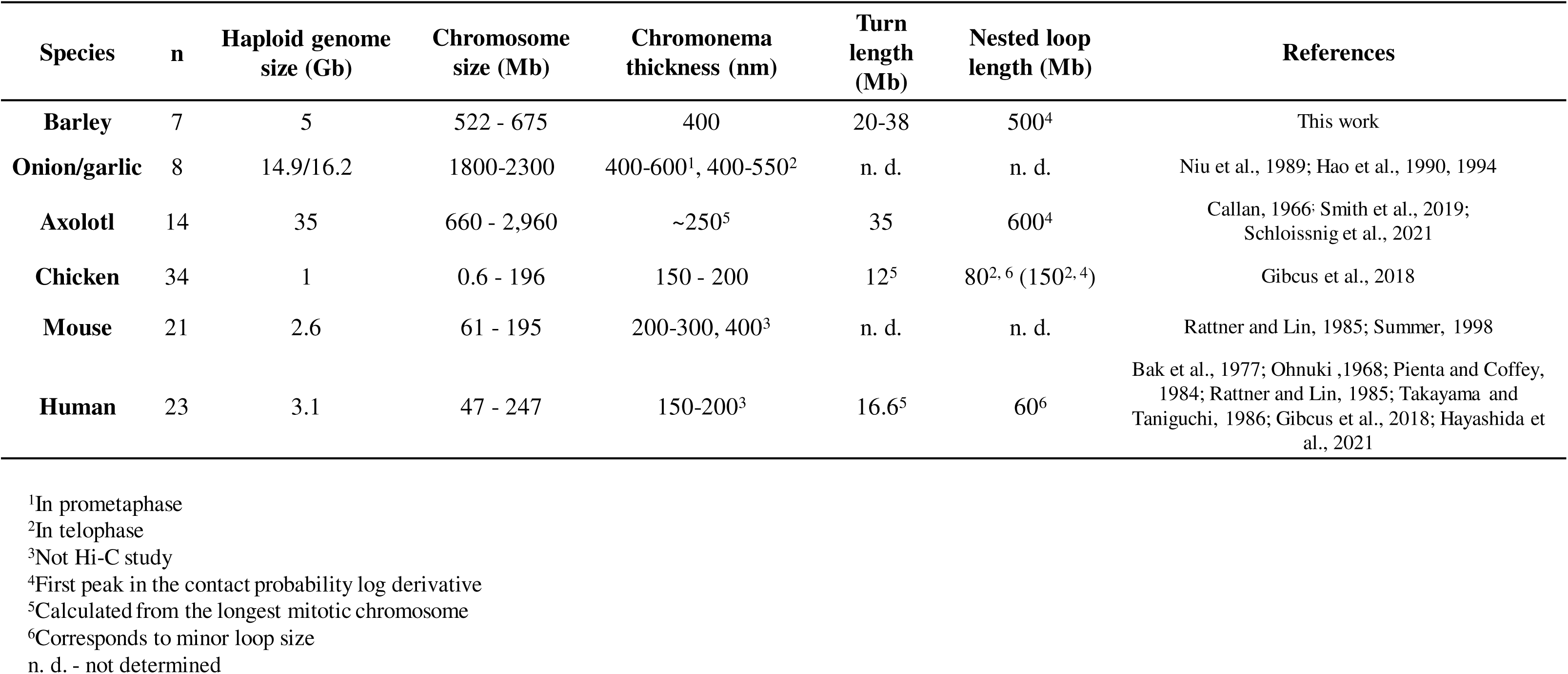
Parameters of helically organized somatic metaphase chromosomes of barley compared to other species. Chromatid helix turn sizes and nested loop sizes are based on Hi-C data and a bottle brush-like model except for mouse and human.

| Species | n | Haploid genome size (Gb) | Chromosome size (Mb) | Chromonema thickness (nm) | Turn length (Mb) | Nested loop length (Mb) | References |
| --- | --- | --- | --- | --- | --- | --- | --- |
| Barley | 7 | 5 | 522 - 675 | 400 | 20-38 | 500<sup>4</sup> | This work |
| Onion/garlic | 8 | 14.9/16.2 | 1800-2300 | 400-600<sup>1</sup>, 400-550<sup>2</sup> | n. d. | n. d. | Niu et al., 1989; Hao et al., 1990, 1994 |
| Axolotl | 14 | 35 | 660 - 2,960 | ~250<sup>5</sup> | 35 | 600<sup>4</sup> | Callan, 1966; Smith et al., 2019; Schloissnig et al., 2021 |
| Chicken | 34 | 1 | 0.6 - 196 | 150 - 200 | 12<sup>5</sup> | 80<sup>2, 6</sup> (150<sup>2, 4</sup>) | Gibcus et al., 2018 |
| Mouse | 21 | 2.6 | 61 - 195 | 200-300, 400<sup>3</sup> | n. d. | n. d. | Rattner and Lin, 1985; Summer, 1998 |
| Human | 23 | 3.1 | 47 - 247 | 150-200<sup>3</sup> | 16.6<sup>5</sup> | 60<sup>6</sup> | Bak et al., 1977; Ohnuki ,1968; Pienta and Coffey, 1984; Rattner and Lin, 1985; Takayama and Taniguchi, 1986; Gibcus et al., 2018; Hayashida et al., 2021 |

<sup>1</sup>In prometaphase <sup>2</sup>In telophase <sup>3</sup>Not Hi-C study <sup>4</sup>First peak in the contact probability log derivative <sup>5</sup>Calculated from the longest mitotic chromosome <sup>6</sup>Corresponds to minor loop size n. d. - not determined

The differences in chromatin conformation between interphase and metaphase chromosomes were also reflected by the relationship of the Hi-C contact probabilities of loci pairs and their distance in the linear genome (d) (Figures 1B, S2). The contact probabilities in interphase showed a uniform exponential decay (∼d-1) from 100 kb to 100 Mb (Figures 1B, S2A, B, E). This pattern is reminiscent of the Rabl conformation observed in anaphase.

In metaphase, the distance-dependent-decay rates were more variable, and its derivative showed two regions with increased contacts, ∼500 kb and 30 Mb (Figure S2C, D, E). Because mitotic chromosomes likely become compacted by chromatin loops (Gibcus et al. 2018), we suggest that the broader peak at ∼500 kb arises from an arrangement of nested loops, where the largest loops (∼3 Mb) appear only in metaphase, as indicated by the first peak in the contact probability ratio between metaphase and interphase (Figure S2F). The highest peak in this contact probability ratio (∼30 Mb), the only one with a positive derivative, corresponds to the second diagonal in the contact matrix (Figure 1B).

Gibcus et al. (2018) proposed that this pattern reflects the arrangement of the chromatids into a helix, whose turn length corresponds to the position of the “bump” in Figure 1B. The arrangement into helical turns is schematically presented in Figure 1C, where the spatial distance between genomic loci is proportional to the contact probability. This observation suggested that the helical coiling could be observed along the chromosome axis when differentially labeled (Figure 1D). We estimated that depending on its size, a condensed barley chromosome consists of 18-23 helical turns (Figure 1E, Table S1).

A closer inspection of the metaphase contact matrix suggested that the distance between the main and secondary diagonal is variable. We analyzed turn sizes along the chromosome lengths based on Hi-C using a sliding window approach and considering only contacts spanning a moving focal point. We observed different local patterns. The characteristic bump of the contact probability shifted (Figures S3, S4; Movie 1) as the focal point moved along the chromosome. The different placements of the bump indicate that the turn length varies along the length of the chromosome. The turn size decreased smoothly, but non-linearly, from the centromere towards the telomere, with turn sizes ranging from 20 to 38 Mb. Moreover, the turn length decreased sharply towards the primary and secondary chromosome constrictions (Figure 1E).

### Oligo-FISH confirms the helical organization of mitotic chromatids

To validate the results obtained by Hi-C, we performed FISH with pooled oligonucleotide (oligo) probes. A search for unique sequences suitable for probe design by scanning Morex v1 (Beier et al. 2017; Mascher et al. 2017) with a 40 Mb window revealed a continuous region of ∼157 Mb with a high density of single-copy sequences on the long arm of chromosome 5H (Figures 2A, S5). We designed FISH probes comprising pools of ∼45 bp oligos to label selected parts of this region; the pooled probes were named Stork, Eagle, Ostrich, Rhea, Moa, and Flamingo. Examples of Stork and Moa probes are shown in Figure S6. Based on the Hi-C data and Morex v1, Stork and Eagle covered half helical turns, whereas Ostrich, Rhea, and Moa each represented one full helical turn. The Flamingo probe was located at the chromosome terminus, just before the subtelomeric and telomeric repeats (Figure 2B, C). The density of single-copy sequences increased towards the telomere (Figure S7), in agreement with the reported higher density of genic and single-copy sequences in distal regions (Mascher et al. 2017).

**Figure 2.**
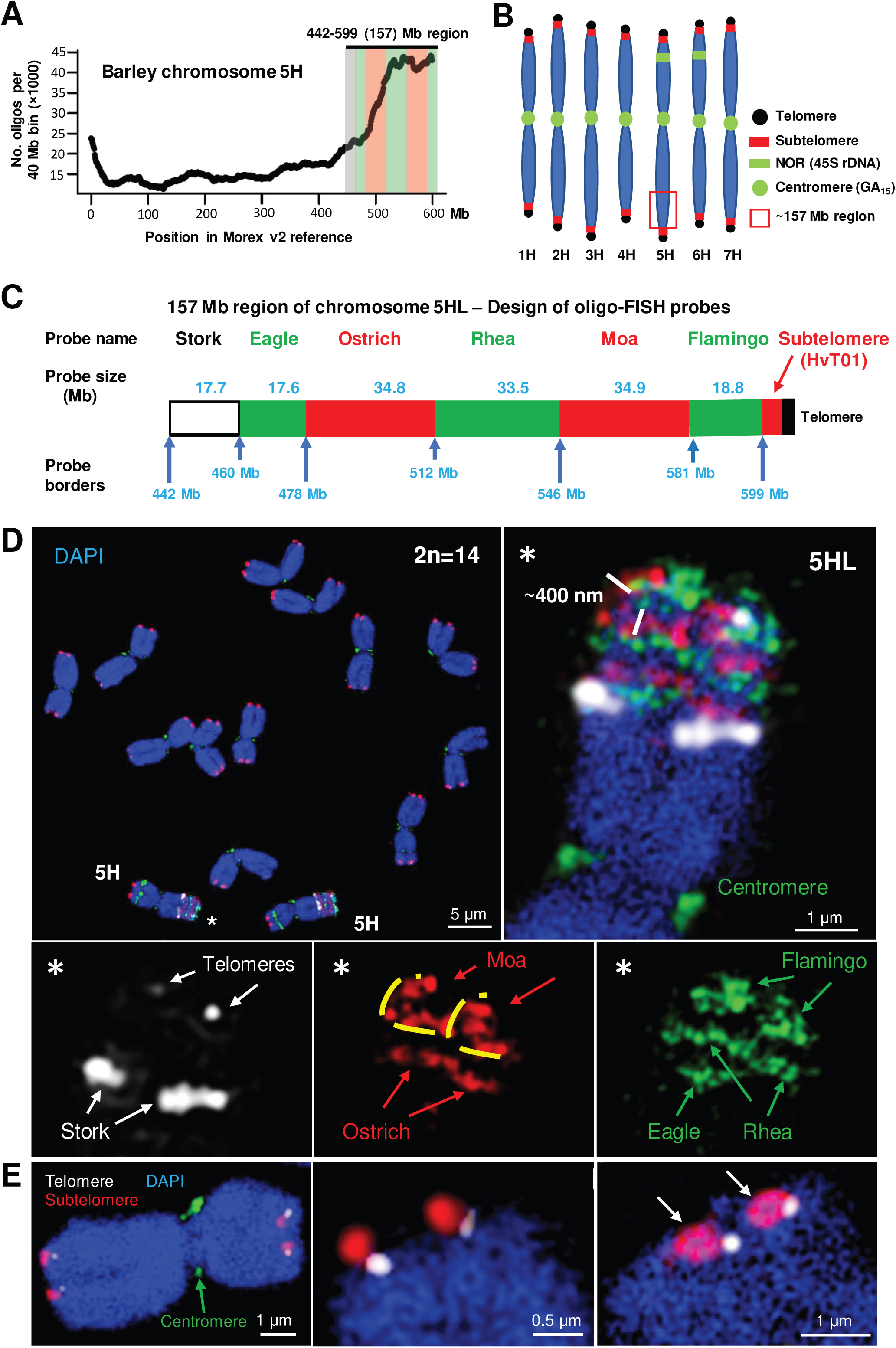
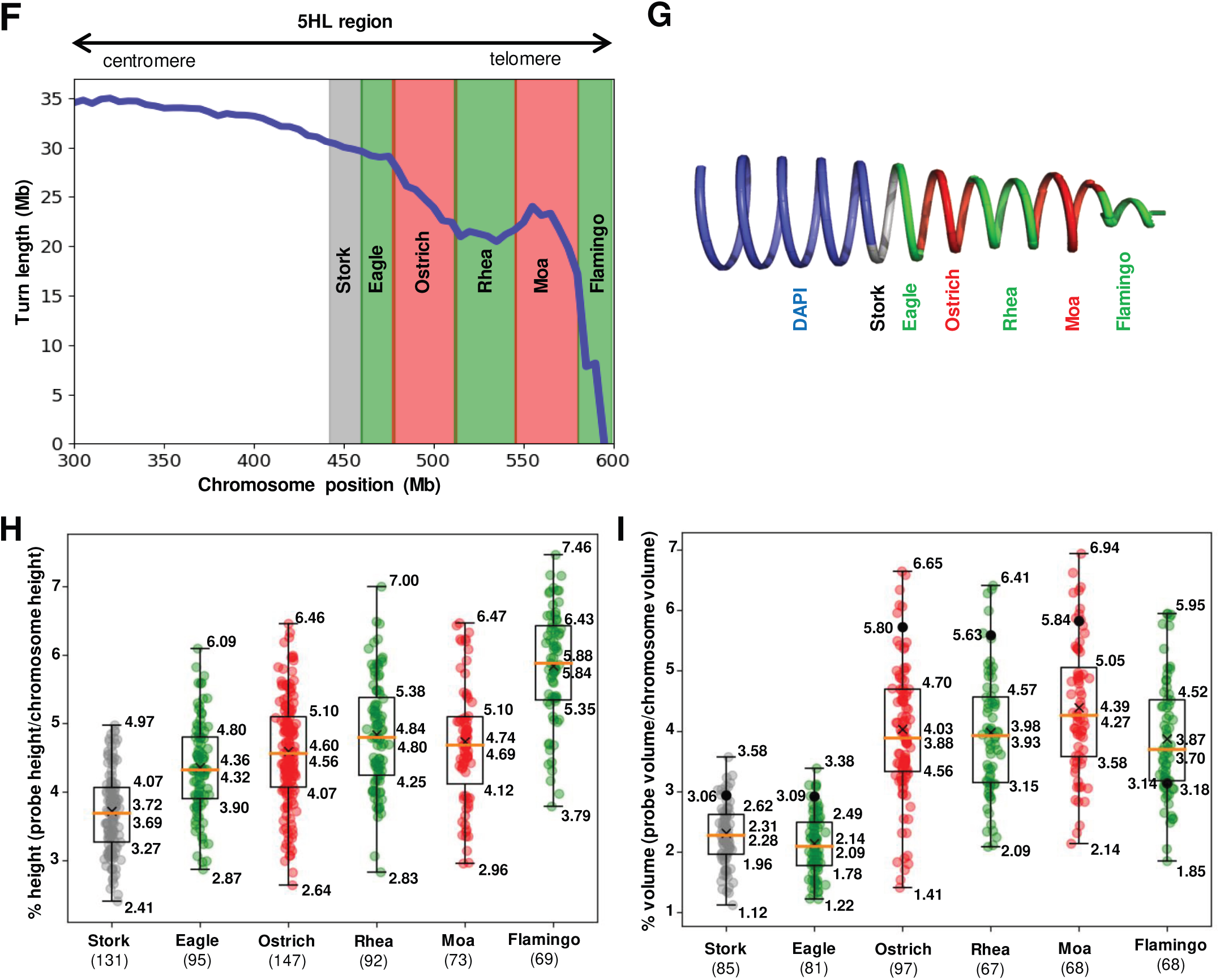
FISH using oligo probes reveal the helical organization of barley metaphase chromosomes. **(A)** Density of predicted single-copy oligo probes along chromosome 5H. The distal 157 Mb-long region of the long arm of chromosome 5H is enriched in single-copy sequences suitable for the detection by oligo-FISH. **(B)** Karyogram of barley indicating the distribution of applied centromere-, NOR-, subtelomere-, and telomere-specific FISH probes. **(C)** Design of the oligo-FISH probes (named after different birds for simplicity) covering the 157 Mb-long region of chromosome 5HL. Precise probe coordinates are given in Table S2. Stork and Eagle together cover a ∼1.2 turn, and Ostrich, Rhea, and Moa cover a ∼1.5 turn. Flamingo covers the subterminal part of the long arm. **(D)** Metaphase chromosome spread labeled with the FISH probes shown in **(B)**. The enlarged oligo-painted region of 5HL (asterisks) shows the chromatin arrangement in both sister chromatids as predicted by the Hi-C-based helical chromatin arrangement model (Figure 1D). Due to chromosome tilting, Moa (in a top-side view) shows the spiral organization of the ∼430 nm thick chromatin fiber (marked with yellow lines). **(E)** Positions of telomeres and subtelomeres vary at both termini of different chromosomes. Subtelomeres form ring-like structures (right, arrows). Chromatin was counterstained with DAPI (blue). **(F)** Turn lengths (blue line) across the 300 Mb region of 5HL encompassing the designed oligo-FISH probes calculated from the Hi-C data. The regions covered by each probe are colored according to the probe color. **(G)** Helical arrangement of the target region illustrating the changes in the turn lengths according to the region shown in (F). The model is compatible with a helical arrangement with similar turn heights and volumes (see H) and with varying turn lengths. **(H, I)** Box-plots showing the relative measured signal heights **(H)** and volumes **(I)** in percent per whole chromosome of the oligo-labeled half (Stork and Eagle)- and full-turn regions (absolute values are visualized in Figure S9). The numbers of measured chromosomes per oligo probe are shown in parentheses. The boxplots show the mean/median (cross/yellow line), lower, and upper quartiles with explicit values. Whiskers extend to 150% of the interquartile region below and above the box (Williamson et al. 1989). The high variability within each probe is due to difficulties in signal detection. The black dots in the right diagram **(I)** are the expected chromosome volume (%) for each probe if the chromatin density were uniform along the entire chromosome. For all probes, this volume is higher than the observed mean, reflecting an underestimation in the measurements. The exception is the Flamingo probe, the only one with a higher observed than expected mean volume. This indicates that the Flamingo region is less dense than the other measured chromosome regions. The signal heights were measured within SIM image stacks using ZENBlack software, and the volumes (see Figure 6C) were measured using the surface rendering tool of Imaris 9.6.

After our probe design, an improved assembly of the barley genome (Morex v2) was published (Monat et al. 2019), resulting in a change in the probe coordinates (Table S2). This had no noticeable effect on the oligo probe distribution or specificity, but the lengths of the labeled intervals changed.

The completeness of the assayed interval, located between 442 and 599 Mb in the Morex v2 5H pseudomolecule, was validated by examining a genome-wide optical map of barley cv. Morex (Mascher et al. 2021). The interval was covered by three optical map contigs (1, 55, and 17), which were concordant with the sequence (Figure S8). A single discrepancy within the interval was a missing sequence of ∼403 kb around position 491 Mb, detected as unaligned ends of the optical map contigs 1 and 55. This indicates that the correct length of the DNA segment labeled by the Ostrich probe is 34.8 Mb.

Three-dimensional structured illumination microscopy (3D-SIM) of chromosomes after FISH with pooled oligo probes (Figure 2D; Movies 2, 3) showed that the chromosome diameter decreased towards the termini. This corresponded with the changes in turn size along the chromosome observed by Hi-C analysis. Once the Morex v2 assembly became available, we re-estimated the turn coverage of the probes. Stork and Eagle cover ∼0.6 of a turn (in a region with a turn size of 29.6 Mb), Ostrich, Rhea, and Moa cover ∼1.5 turns (in a region with a turn size of 22.5 Mb), and Flamingo covers a region of ∼2.3 decreasing turns (in a region with a turn size of 8.3 Mb). Thus, the probes cover fewer turns in regions with larger turn sizes (Figure 2F, G; Table S2).

The signals of the oligo-FISH probes Stork and Eagle showed an average height of 380 nm. Although three FISH probes (Ostrich, Rhea, and Moa) represented more than one full helical turn and were labeled by fluorescence dyes delivering a different SIM resolution (laterally ∼120 nm for Alexa488 and ∼140 nm Atto594), the signals of all three probes had similar dimensions (Figures 2H, S9A). On average, these signals were ∼460-480 nm in height. As Hi-C predicted 21 turns for chromosome 5H (Figure 1E), this corresponds to a chromosome length of 9.0–9.4 µm, which is consistent with our chromosome length measurements (Figure S9A).

Based on these observations, we call the helically arranged 400-nm chromatin fiber chromonema. A partial intermingling of smaller chromatin loops inside the chromonema was identifiable between neighboring turns of the helix (Figures 2D, S10; Movie 4). Therefore, the chromatin was arranged in lower-order loops within the chromonema, with the intertwining possibly caused by the dynamic properties of the loops. Remarkably, the orientation of the FISH signals was always identical in both chromatids, indicating that both chromonemata were turned with the same handedness (Figure S11).

We performed surface rendering of the 3D-SIM image stacks and determined the volumes of the different oligo-FISH signals and the whole chromosome 5H (Figures 2I, 6C, S9B; Movie 5). To avoid artificial flatting of chromosomes after preparation, we embedded the isolated chromosomes in polyacrylamide and imaged them by 3D-SIM after DAPI staining. These spatially preserved chromosomes showed ∼1.2-fold higher volumes than chromosomes that were dried on slides for subsequent FISH (Table S3). The volume of an embedded 5H chromatid (∼599 Mb) was ∼24.75 µm3, corresponding to ∼24.2 Mb/µm3, i.e., 1 Mb chromatin is folded into ∼0.041 µm3. In the Flamingo-positive chromosome region, the Hi-C data indicated a smaller turn contact pattern (Figure 2F, G). The volume of the Flamingo signal, the only one larger than expected for a uniformly dense chromosome (Figure 2I), suggested a lower chromatin density (DNA amount per µm3). Compared to Flamingo (19.5 Mb/µm3), the regions of the Stork (32.0 Mb/µm3), Eagle (34.3 Mb/µm3), Ostrich (38.2 Mb/µm3), Rhea (34.0 Mb/µm3), and Moa (32.0 Mb/µm3) probes were ∼1.7 times more condensed. Thus, the terminal Flamingo region was characterized by lower chromatin density and smaller turn size. The flexibility of the chromatin arrangement at the chromosome ends was also visible after FISH with subtelomeric and telomeric probes. Their positions varied at the chromosomal termini, and the subtelomeres showed ring-like shapes, as described by Schubert et al. (2016a) (Figure 2E; Movie 6).

In short, these cytological observations suggest that both metaphase sister chromatids are formed by a ∼400 nm thick chromatin fiber arranged as a helix of identical handedness with decreasing turn size towards the telomeres. The chromosome ends are less compact, possibly allowing for higher chromatin fiber flexibility than in interstitial arm regions.

### Analysis of SCEs confirms the helical 400 nm chromonema structure

The harlequin staining pattern after differential labeling of sister chromatids with base analogs is caused by SCEs and allows chromatin segment exchanges even smaller than the width of a chromatid to be detected. Based on this observation, metaphase chromatids were previously proposed to be formed by a chromatin helix in onion (Schvartzman and Cortés 1977; Schvartzman et al. 1978). We applied the same approach to confirm the spatial chromatin organization of mitotic barley chromosomes.

The extension of the exchanged chromatid segments differed in height and width. The measurements were done parallel and perpendicular to the chromatid axis (Figure 3). Although both dimensions were affected by the degree of chromosome condensation, all exchanged regions shared the following patterns. The height never dropped below 300 nm, the minimum size that may be exchanged. The maximum height was restricted by the chromosome arm length. Importantly, the height of the exchanged part never exceeded 300–550 nm before the exchanged part spanned the full chromatid width (Figure 3B). These findings agree with the helical chromatin arrangement determined by Hi-C and oligo-FISH.

**Figure 3.**
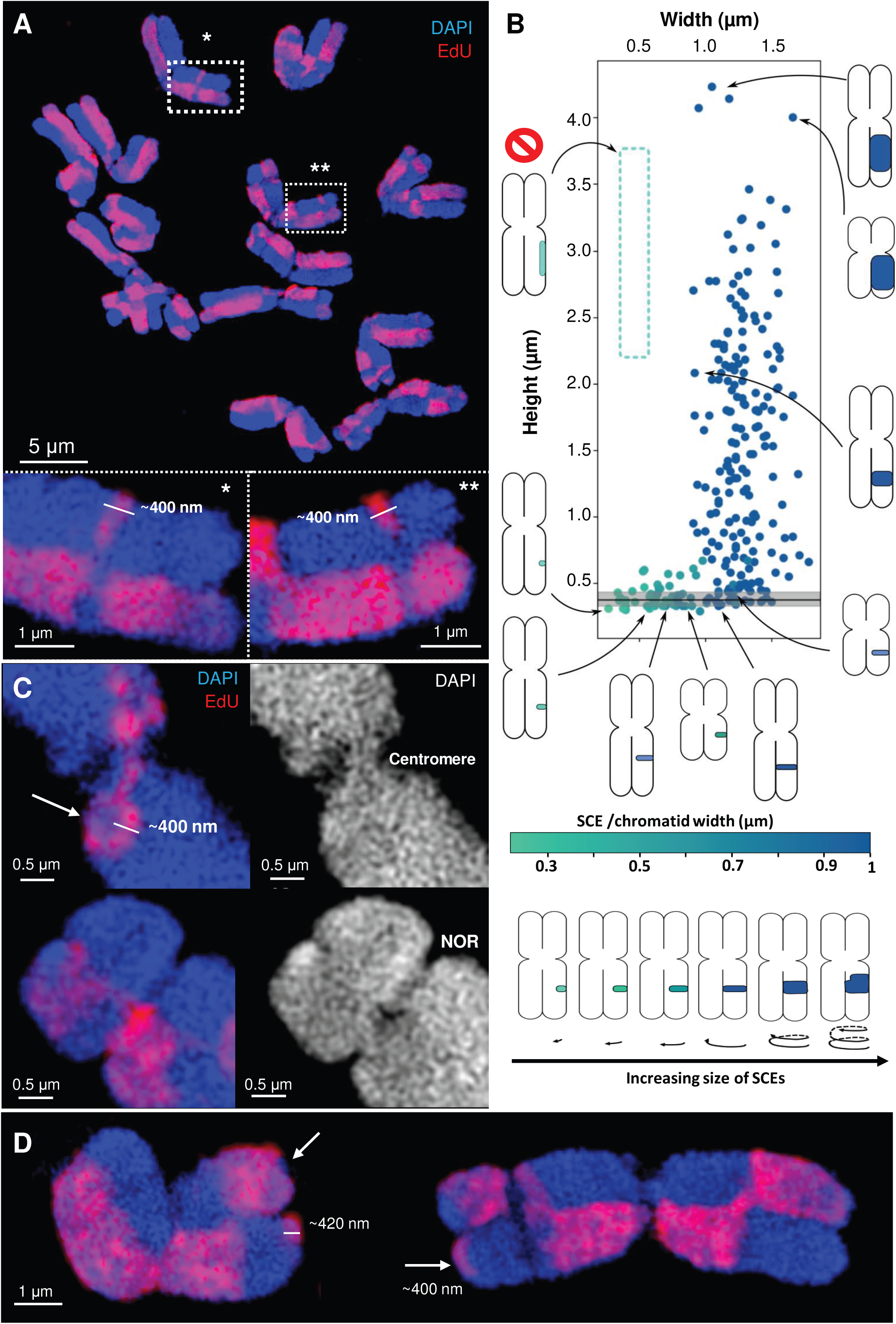
Sister chromatid exchanges (SCEs) confirm the helical organization of barley metaphase chromosomes. **(A)** Metaphase chromosome spread labeled with EdU to detect SCEs creating a harlequin pattern. Some exchanged chromatin regions appear as thin bands with a minimum height of ∼400 nm (enlarged regions * and ** in dashed rectangles). The sizes of these regions correspond to the ∼430 nm chromatin fiber (chromonema) detected by the oligo-FISH probes (Figure 2D, H, S9A), reflecting a helix within the chromatid arms. **(B)** Scatter plot of 257 measured SCEs of different heights (along the chromosome longitudinal axis) and widths (in the chromatid). The color code indicates how much of the chromatid width is covered by the SCE: dark green indicates complete coverage. The schematic chromosomes around the diagram show examples of measured SCEs. The lack of values in the dashed region indicates that SCEs > ∼400 nm and not covering the complete chromatid width never occurred. The black line indicates the median height (∼380 nm) of SCEs with incomplete width (less than chromatid width), and the gray area spans the lower and upper quartiles. The observed possible configurations of SCEs imply that the chromatids are formed by ∼400 nm high chromatin units. These configurations are compatible with a helical arrangement, as illustrated in the scheme below. The small SCEs in the centromeric, NOR and telomeric regions were not considered. **(C)** At centromeres (top), the EdU labeling indicates the presence of a straight, thin connection between the ∼400 nm high fibers. Immediately next to the centromere, due to chromatin tilting, a ∼400 nm thick turn is visible in the pericentromeric region (arrow). At the NOR (bottom), thin chromatin fibers also occur. **(D)** SCE units of ∼400 nm are also evident at the chromosomal termini (arrows). Chromatin was counterstained by DAPI (blue/white).

We then combined SCE detection and FISH with the Stork, Eagle, Rhea, and Flamingo oligo- probes to relate the exchanged chromatid segments to the FISH-painted regions. As shown in Figure 4 and Movies 7 and 8, both 5H homologs contained a small SCE segment close to a Rhea- positive region, and the adjacent regions not painted by FISH with the Moa and Ostrich probes. The shapes and heights (∼400 nm) of the exchanged chromatid segments and FISH signals were similar and partially replaced each other. These findings further confirm the helical chromatin arrangement of metaphase chromatids.

**Figure 4.**
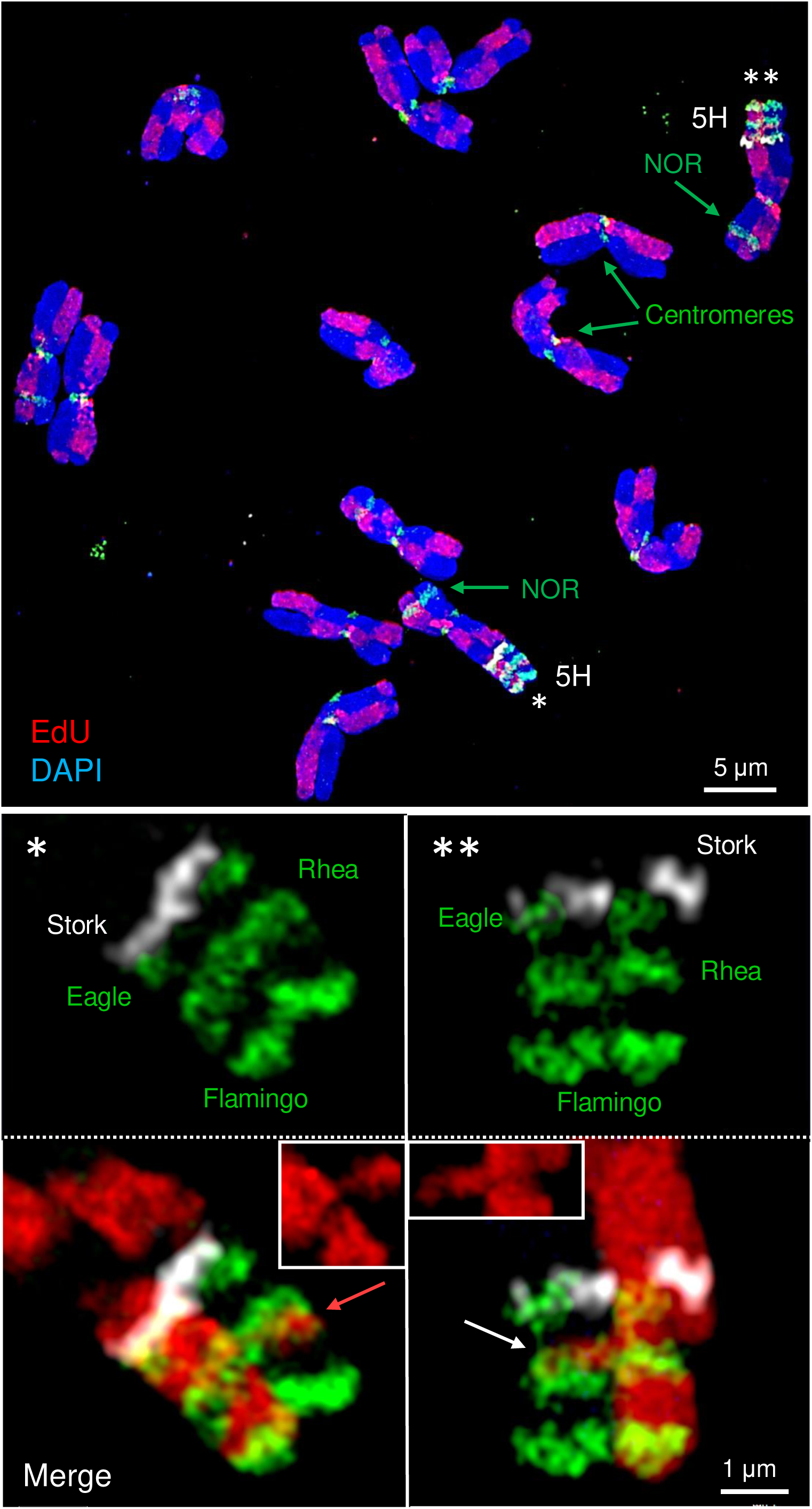
A combination of oligo-FISH and EdU labeled SCEs support the helical organization of chromatin. Metaphase chromosome spread (top) showing all barley chromosomes after EdU and oligo-FISH labeling. In both homologous oligo-FISH-painted 5HL regions, two thin exchanged regions occurred (bottom). The one on the left does not span the entire chromatid width, but the one on the right does (insets in the merged images). The SCEs are present at the transition from the Rhea region to the unlabeled Moa (left, red arrow) and Ostrich (right, white arrow) regions. The heights of the SCEs, oligo-FISH probe and EdU-free regions are similar (∼380-450 nm).

In contrast to the remaining chromatids, no helical organization was identified in the centromeric regions and secondary constrictions (NORs). Instead, straight chromatin fibers were observed (Figure 3C).

### The 400-nm chromonema consists of loops of ∼80-nm chromatin fibers and shows no large cavities

In addition to the ongoing discussion about the helical versus non-helical higher-order metaphase chromosome structure, it remained an open question whether the chromatin-free ‘cavities’ along the entire lengths of chromatids detected in several electron microscopy investigations in plants and human (Hao et al. 1988, 1990; Hamano et al. 2014; Kuznetsova et al. 2017) reflect a real phenomenon or simply a preparation artifact. Thus, we stained barley chromosomes with DAPI to reach a 3D-SIM resolution of ∼130 nm (Kubalová et al. 2021) for reliable observation of the spatial ultrastructure (Figure 5). Large cavities were never identified within complete 3D-SIM image stacks of metaphase chromosomes. Instead, we detected a network of looped ∼80-nm chromatin fibers (n=36; mean 77.2±7.7). Between these fibers, chromatin-free regions no larger than ∼120×220 nm were evident (Movies 9, 10). The 80-nm fibers formed the 400-nm chromonemata. At centromeres and NORs, both looped and straight ∼80-nm fibers were observed. These fibers apparently represent a constriction-specific substructure of the chromonema which is not helically arranged in these regions.

**Figure 5.**
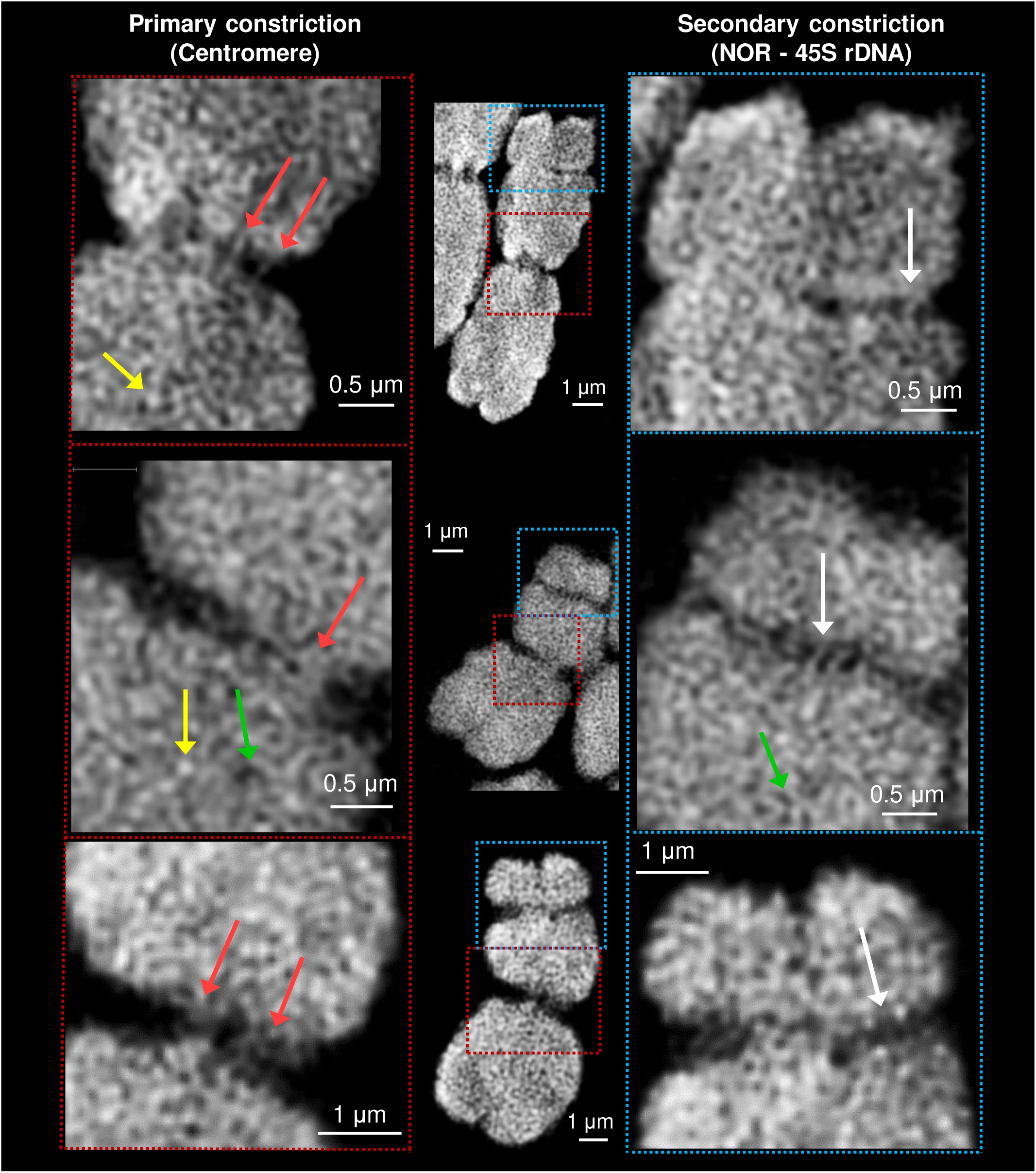
Primary (centromere) and secondary (NORs) chromosome constrictions are not organized as a helix. Instead, these regions (marked by red and blue dashed rectangles) contain parallel chromatin fibers (red and white arrows; also see Figure 3C). At chromosome arms, the DAPI-labeled chromatin (white) occurs as a network of looped fibers within the ∼400 nm fiber. The fibers contain chromatin clusters (yellow arrows), and in between them, small chromatin-free regions (green arrows). No large continuous cavities were observed, as demonstrated in Movies 9 and 10, representing the whole chromosome volume.

### Polymer modeling confirms the helical chromatid architecture

We constructed polymer models using the calculated turn sizes to reproduce the bump position in the contact probabilities of mitotic chromosome Hi-C data and the arrangement of the chromosome regions highlighted by oligo-FISH (Figure 6). The models were built with equilibrium molecular dynamics simulation based on the bottle-brush model proposed by Gibcus et al. (2018). The models were created by large side-by-side loops, whose bases followed a helical path. The large loops consisted of minor side-by-side loops that filled the space surrounding the helical path, both internal and external to it. The loops formed turns of chromatin that stacked along the chromatid axis (Figure S12D).

**Figure 6.**
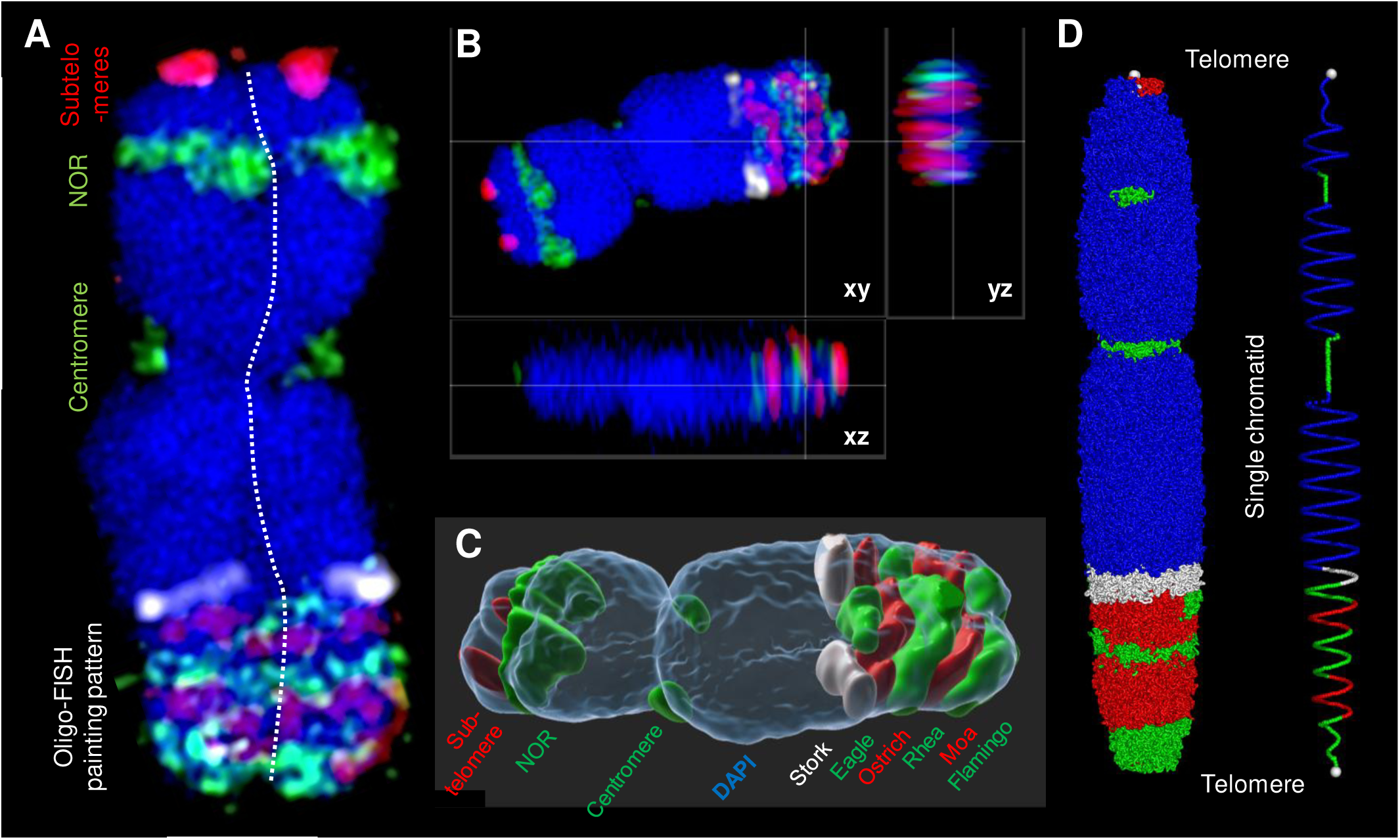
Simulated Hi-C data-based polymer chromosome model based on cytological findings. **(A)** Metaphase chromosome 5H labeled by different FISH probes (marked by an asterisk in Figure 2D). The chromatids are separated by a dashed line. The ortho-view (Movie 3) **(B)** and surface rendering (Movie 5) **(C)** of the same chromosome prove that the FISH signals are present throughout the whole chromosome volume. **(D)** A polymer model of a single chromatid based on the turn lengths calculated from the experimental Hi-C data (Figure S14). The simulated model, built as a bottle brush, is formed by large loops, whose bases follow a helical arrangement (right). The large loops are further divided into minor loops, which internally and externally fill the space surrounding the helical arrangement. Thus arranged, the loops form thicker helical turns of chromatin. Each monomer in the model represents ten nucleosomes (∼2 kb). The entire polymer contains 300,000 monomers corresponding to ∼600 Mb. The multi-colored region follows the design of the oligo-FISH probes (Figure 2C). The model can be used to illustrate the number and changing sizes of the turns along the chromatid arms, but it has to be adapted according to the observed FISH-labeling pattern, e.g., the NOR appears larger via FISH labeling than that simulated based on the Hi-C data.

**Figure 7.**
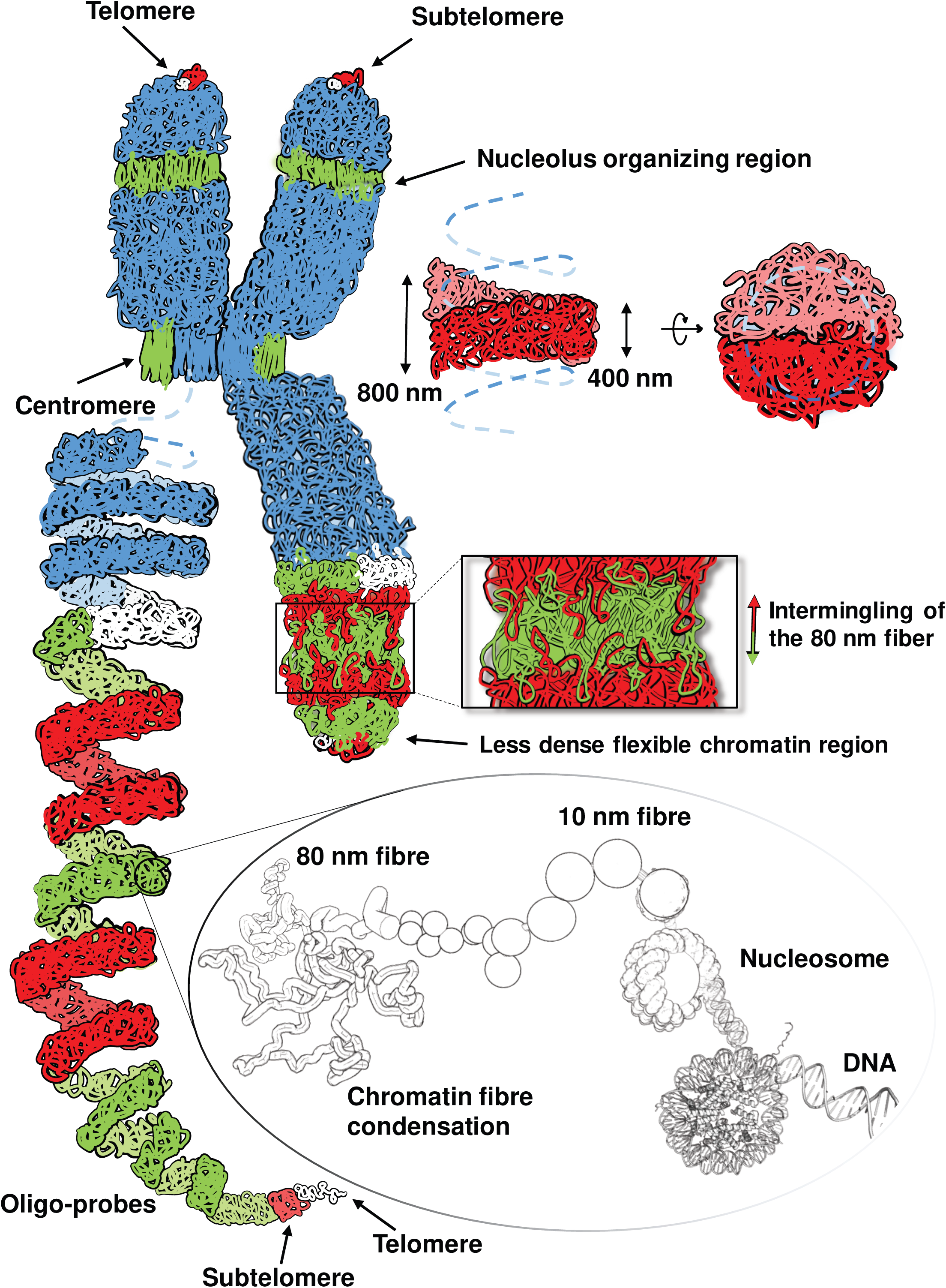
Helical model of the higher-order chromatid organization of a somatic plant metaphase chromosome based on data from barley. Both chromatids (the highest level of chromatin organization) are formed by a ∼400 nm thick chromatin fiber (chromonema) arranged in a helix of identical handedness (Figure S11). The chromatids are filled with chromatin without large cavities. Adjacent chromonema turns intermingle due to the flexibility of the smaller 80 nm fibers present within the chromonema. The helical order is interrupted at centromeres and secondary constrictions, which mainly display straight 80-nm chromatin fibers. The chromosomal termini contain less condensed, more flexible chromatin. Due to this flexibility, the telomeres may be embedded into the subtelomeric chromatin and not appear at the very end of the chromatid. The left long arm chromatid is shown as a stretched helix representing their higher-order chromatin folding based on oligo-FISH labeling. The different colors represent half (white + green) and complete (red + green) chromonema turns, respectively. The chromonema is formed by looped, thinner 80 nm chromatin fibers. This lower- order chromatin arrangement (colorless inset) is still under debate. According to previous findings (^9^Thoma et al. 1979), at the lowest hierarchical level, 147 bp of left-handed DNA is wrapped around a nucleosome core. Nucleosomes with DNA linkers in between them form the 10-nm chromatin fiber. Several nucleosomes associate into thicker 80 nm chromatin fibers, which form loops within the chromonema.

To compare the polymer models with the Hi-C contact probability data, we tested whether our simulation reproduced the contact probability of chicken chromosome 1 at prometaphase, as described by Gibcus et al. (2018) (Figure S12). We simulated only five turns of the helical arrangement (with constant turn lengths) using a model of the 10-nm chromatin fiber, where each monomer represents one nucleosome.

The simulated model for chicken closely followed the experimental contact probability. To reproduce the bump that we observed in the Hi-C contact probabilities of the barley chromosomes, we set the turn length to 30 Mb and the turn height to 400 nm. However, our barley models 1 –4 did not reproduce the contact probability as well as our model using the data from chicken chromosomes (Figure S12A-C). Therefore, we considered model 5 where the chromatin loops were larger, leading to a dispersed helical protein scaffold (Figure S12E). This chromatin loop arrangement followed the Hi-C data, suggesting (on average) large 3-Mb loops and small 500-kb loops (Figure S2E, F). Accordingly, we suspect that the loop arrangement differs in barley compared to other species with possible differences in the density and size of loops surrounding the helical path.

To consider a half-helical arrangement of chromatids as suggested by Chu et al. (2020), we simulated a 10-Mb region (to reduce computational costs) with coarse-grained models of the 10- nm chromatin fiber. In this model, the helical path changes the turn handedness (left to right) every half turn. The turn length was set to 2 Mb (making 10 half-turns), but instead of a lower contact probability at the half-turn distance (1 Mb), which is characteristic of helical arrangements, we observed a plateau. Sites separated in the genome sequence by half a turn length may be spatially far or close, but in the helical arrangement, they are always far apart. The irregularity of half-turn contacts broadens the bump for inter-turn contacts. We expect that other irregular structural features, such as turns with different lengths and perversions between half-turns, as suggested by Chu et al. (2020), would only broaden and not sharpen the contact probability peak for distances in the range of a turn length (Figure S13).

To achieve a model matching our cytological observations, we simulated a 600 Mb chromatid with a coarse-grained model, where each monomer represents 2 kb (Figure 6D; Movie 11). The bases of the loops were constricted to a helical arrangement, with turn lengths as calculated from Hi-C data (Figure 1E). The resulting model for chromosome 5H has seven turns in the short arm and 14 in the long arm, which fits with the estimation derived from cytogenetic observations. Regions labeled by oligo-FISH and representing successive helical turns appear as stacked layers, consistent with our cytological observations. The Stork and Eagle oligo probes (encompassing less than one turn) present signals with lower heights (corresponding to one-turn height) and do not fully occupy the width of the chromatid (depending on the viewpoint). Signals using Ostrich, Rhea, Moa, and Flaming oligo probes (encompassing more than one turn) are taller and always occupy the entire width of the chromatid. The variable turn lengths in the model reproduce the cylindrical shape of the chromosome, with a decreasing radius towards telomeres and constrictions. Replicates with different and randomly chosen loop sizes (see Materials and Methods) introduced variability into the model, where the labeled regions have slightly different shapes (Figure S14). The loop variability and flexibility should account for the expected heterogeneity of the cell population used in the Hi-C experiment (Shi and Thirumalai 2021), while the underlying main helical organization holds even for individual chromosomes, as shown by our cytological results.

Other experimentally observed features could not be reproduced by the simulated model. Subtelomeric and telomeric regions, which were mainly absent from the genome assembly, may be less dense than simulated, as indicated by volume measurements (Figures 2I, S9B). Sequences of the primary and secondary constrictions were also lacking in the genome assembly, and the less clear organization of centromeres and NORs precludes modeling of their conformations.

## Discussion

### Helical metaphase chromosome organization – a conserved feature?

A helical structure of plant chromosomes was previously seen in living tissues but also after physical treatments by a wide range of agents on fixed chromosomes. The coiled chromatin fiber forming chromatids (called chromonema in early publications) has been observed in plants from different families during mitosis and meiosis. The coiling of mitotic chromosomes can vary from loose and irregular to the formation of a regular compact helix. Various authors originally described two chromonemata per chromatid (Nebel 1932a, 1939; Sax 1935; Matsuura, 1935, 1938; Manton 1950). However, Ruch (1950) clarified in *Tradescantia virginica* that these observations were based on optical artifacts and postulated that only one chromatin helix was present per chromatid. Our investigations based on molecular, cytological, and super-resolution microscopy data provide independent evidence that one helically coiled 400-nm chromatin fiber forms a metaphase chromatid in barley, corresponding to the classically named chromonema in plants. Similarly, a helical chromatid organization was visualized by electron microscopy in large-genome plants including onion (Hao et al. 1990), barley (Iwano et al. 1997), and faba bean (Döbel and Schubert 1981). A helical structure was also observed in chromosomes of animals such as frog (Seto 1972), axolotl (Schloissnig et al. 2021), mouse (Sumner 1998; Rattner and Lin 1985), Chinese hamster (Ris 1981), and human (Ohnuki 1965, 1968; Utsumi 1982; Rattner and Lin 1985; Boy de la Tour and Laemmli 1988; Sumner 1991, 1998; Phengchat et al. 2019; Hayashida et al. 2021).

The available data suggest that the longer and thicker chromosomes of species with larger genomes contain more and larger chromonema turns and longer inside loops (Table 1). In human, the number of turns is positively related to chromosome length (Boy de la Tour and Laemmli 1988). If we consider the 15 turns Ohnuki (1968) observed in chromosome 1 and the chromosome size of 248.96 Mb (*Homo sapiens* GRCh38.p13 genome assembly), the average size of one turn should be 16.6 Mb. This is similar to the sizes (12 Mb and 10 Mb) determined by Gibcus et al. (2018) using Hi-C for the helical turns in chicken and human chromosomes, respectively. In contrast to the macrochromosomes of chicken, the chromatids of chicken microchromosomes, which are smaller than 12 Mb, do not form turns (Gibcus et al. 2018). The diameter of the helically coiled chromatin fiber forming mouse chromatids was estimated to be 200-300 nm (Rattner and Lin 1985). A single turn of the helically organized 28 mitotic chromosomes of the giant axolotl genome (32 Gb/1C) comprises ∼35 Mb DNA (Schloissnig et al. 2021).

The number of turns was reported to vary from 16 to 46 in mitotic metaphase chromosomes and from 3 to 23 in the more condensed meiotic metaphase chromosomes in plants of the genera *Fritillaria, Osmunda, Rhoeo, Sagittaria*, *Tradescantia*, and *Trillium* (Manton 1950). We determined that each barley chromosome (522 to 675 Mb long) contains 18-23 turns of 20-38 Mb along the arms, depending on chromosome length and position. Towards the chromosomal termini, the turns become smaller and the chromatin is less densely packed. The different positions of subtelomeres and telomeres in both sister chromatids point to the flexibility of the lower-order chromatin structural organization. Using FISH on longitudinally stretched barley chromosome 1H, Valárik et al. (2004) found that telomeric sequences do not localize to the very ends of the chromosome. Ring-like subtelomeric repeat arrangements (Schubert et al., 2016a) may reflect the helical higher-order chromatin structure at the chromosome termini. The decondensation of barley telophase chromosomes starts at the chromosome ends, possibly due to lower chromatin density and higher flexibility (Wanner and Formanek 2000).

The chromosome length does not affect the average turn length within a species. We compared the contact probability of condensed chicken (Gibcus et al. 2018) and barley chromosomes, finding that the turn length differs between species but not in the same species (Figure S15). This is more obvious for chicken, whose chromosomes range from 0.6 to 196 Mb long. Only chromosomes smaller than a turn length (12 Mb) lack the characteristic bump in the contact probability diagram. A comparison between the contact probability derivatives of chicken and barley suggested that the chromatin loop arrangement within the chromonema can also differ between species but is similar between the chromosomes of the same species. Therefore, chicken shows smaller chromatin loops than barley.

The second parallel diagonal in the Hi-C contact matrix, and the respective bump in the contact probability, which relate to turn length, are key features explainable only by a helical arrangement of mitotic chromatids (Figure 1). Other arrangements modeled by polymer simulation, such as an equilibrium globule, fractal globule, and linearly compacted loops (Mirny 2011; Naumova et al. 2013), resulted in different contact patterns (Figure S13). Here, we also simulated a half-helical model, where the handedness changes every half turn, similar to the model proposed by Chu et al. (2020). However, in this case, half-turn distances are not always apart in space, and full-turn distances are not always close in space. Thus, this model had to be rejected, as the contact probability lacks the characteristic bump between half-turn and full-turn distances (Figure S13). Using bottle-brush polymer models (Gibcus et al. 2018) and following a helical arrangement, we successfully reproduced the chicken contact probability. Our model for barley mitotic chromosomes, with a turn length of 30 Mb, correctly positioned the bump in the contact probability, confirming that the bump originates from the helical arrangement and is related to turn length. Assuming a height of 400 nm, as detected cytologically for the barley chromonema, we further approximated the model to the experimental observations. By varying the lengths of major and minor loops fitting to the experimental Hi-C data, we induced a dispersed helical scaffold (Figure S12E) and improved the contact probability. The various chromonema helix turn lengths and heights in barley and other species (Table 1) suggest a different mitotic coiling of the major chromatin loops. Due to the different minor loop sizes in these species, the major chromatin loops may attain different widths, resulting in different turn heights. The extent of these differences and how the chromonema coils remain to be elucidated.

Paulson and Laemmli (1977) suggested a ‘scaffold model’ based on histone-depleted human chromosomes, where a non-histone protein backbone is responsible for chromatin looping and chromosome condensation. Topoisomerase IIα and the Structural Maintenance of Chromosomes (SMC) complexes condensin I and II are components of the scaffold in *Drosophila* (^104^Sureka et al., 2018), chicken (Ohta et al. 2019), and human (Maeshima and Laemmli 2003; Fukui and Uchiyama 2007; Maeshima et al. 2018; Walther et al. 2018) and are also evident in *Arabidopsis* and barley (Kubalová et al. 2021; Municio et al. 2021). In human HeLa cells, the chromatid axes appear as isolated compaction centers rather than forming a continuous scaffold (Sun et al. 2018) and appear to consist of a helical structure that serves to organize chromatin loops into the metaphase chromatid (Phengchat et al. 2019). The network-like distribution of topoisomerase IIα that we previously identified in barley metaphase chromosomes (Kubalová et al. 2021) fits the predicted Hi-C data-based dispersed helical scaffold (Figure S12E).

Chromosome compaction by condensin II during mitosis appears to be a conserved mechanism influencing the arrangement of chromosomes in interphase nuclei (Hoencamp et al. 2021), such as the Rabl configuration (Rabl 1885) present in barley (Jasencakova et al. 2001; this work). Further studies on the localization and quantification of scaffold proteins may allow a functional model of condensed barley chromosomes to be established, including lower-level chromatin organization. We propose that the helical arrangement is evolutionarily conserved in large (>12 Mb) metaphase chromosomes of plants and animals.

### The chromatid helix turning direction is random

In the oligo-FISH-labeled regions of both mitotic barley 5H chromatids, we always observed the turning of the chromonema in identical directions. Sister chromatids of human HeLa chromosomes have predominantly opposite (mirrored) helical handedness (Boy de la Tour and Laemmli 1988). Because sister chromatids start to separate at the beginning of prophase (Nagasaka et al. 2016), the turning direction must already be determined after replication during the G2 phase, but it seems to be unrelated to the left-handedness of chromatin fibers, as reported for the salamander *Necturus* (Williams et al. 1986). Based on investigations of *Drosophila* polytene chromosomes, Sorsa (1986) suggested that the size and accumulation of chromomeric loops cause the chromonema to bend and form a helix.

In the plant genera *Trillium*, *Rhoeo*, *Osmunda*, and *Vicia*, the turning direction can change at the centromere and in different regions along the arms, suggesting that no uniform control mechanism functions throughout the whole chromosome. The ratio of left-handed to right-handed turning appears to be random. Only in certain *Tradescantia* genotypes does an excess of right-handed segments point to specific genetic control (Nebel 1932a, b; Sax and Humphrey 1934; Sax 1935; Nebel and Ruttle 1936a, b; Manton 1950; Manton and Smiles 1943). In short, it appears that the higher-order helical turning direction of chromonemata is flexible rather than strictly determined.

### Why is the chromatid helix interrupted by straight and less condensed chromatin at centromeres and NORs?

Using super-resolution microscopy, we detected thinner, straight chromatin fibers at the monocentromeres and secondary constrictions of barley lacking the helical chromonema organization. This finding is in line with the observations achieved by scanning electron, helium ion, and classical light microscopy in barley (Seto 1972; Iwano et al. 1997; Wanner et al. 1991; Wanner and Formanek, 2000; Schroeder-Reiter and Wanner 2009; Schroeder-Reiter et al. 2012; Sartsanga et al. 2021), faba bean (Maruyama 1983; Schroeder-Reiter and Wanner 2009), frog (Seto 1972), Indian muntjac (Rattner 1987), wallaby, and human (Utsumi 1982; Schroeder-Reiter and Wanner 2009). The 200–300 nm chromonema of mouse, consisting of a coiled 100 nm sub-fiber, appears straight in the centromere region (Rattner and Lin 1985, 1987). Even at elongated pea centromeres (Neumann et al. 2012) and along holocentric chromosomes of the grass *Luzula nivea*, parallel chromatin fibers are evident (Schroeder-Reiter and Wanner 2009).

DNA-specific dyes stain primary and secondary constrictions less strongly than chromosome arms, which may reflect a lower chromatin density. Electron microscopy revealed that the amount of looped chromatin fibers and ‘knobby’ chromatin, abundantly visible along the chromosome arms, is reduced in barley centromeres, displaying less DNA but more protein than chromosome arms (Wanner et al. 1991; Iwano et al. 1997; Wanner and Formanek 2000; Schroeder-Reiter and Wanner 2009). Bailey et al. (2013) showed that epigenetic modifications of the N-terminal tail of the centromeric histone variant CENH3 influence the physical properties of chromatin fibers in HeLa cells such that the chromatin organization at centromeres may be altered. Transcription of centromeric DNA during mitosis (Liu et al. 2021) could also contribute to this alteration.

Features of active rDNA might be maintained during mitosis, preventing a helical chromonema structure at NORs. Not only microscopy data but also our Hi-C data revealed a peculiar non-helical chromatin architecture around the NORs and centromeres, manifested as regions without periodical contact patterns in Hi-C contact maps. Perhaps the local loss of long-range contacts is prevented by proteins associated with NORs and centromeres. Data obtained by chromatin conformation capture and analysis of nucleolus-associated chromatin domains in *Arabidopsis thaliana* support this hypothesis for NORs (Feng et al. 2014; Pontvianne et al. 2016) and centromeres (Grob and Grossniklaus 2017). Alternatively, perhaps the missing contacts in the Hi-C matrices are due to the absence of centromeric and rDNA sequences in the reference genome of barley and/or the highly repetitive nature of surrounding regions, precluding unique contact mapping.

Overall, we speculate that the missing helical arrangement and the relaxed chromatin allow kinetochores to be established at centromeres and maintain the structures of transcriptionally active NORs. The identification of several parallel ∼80-nm chromatin fibers within both constrictions suggests a specific arrangement of the chromonema, which remains to be elucidated.

### Lower-level chromatin organization – a network of chromatin globules and looped fibers

Without specific labeling, the tight compaction of metaphase chromosomes makes individual helical turns invisible. The intermingling of lower-order chromatin fibers between adjacent turns of the chromonema in barley (visualized by oligo-FISH) explains this observation. Super-resolution microscopy revealed a network of globular (clustered) and fibrous chromatin structures within and at the surface of barley metaphase chromatids, similar to the fibers and ‘knobby’ structures possibly corresponding to chromomeres (coiled solenoids) (Serra 1947; Zatsepina et al. 1983) observed by scanning electron microscopy at the surfaces of barley (Wanner et al. 1991; Iwano et al. 1997; Wanner and Formanek 2000; Sartsanga et al. 2021), Chinese hamster (Zatsepina et al. 1983), and human chromosomes (Sumner 1991). Because our Hi-C contact matrices from barley inter- and metaphase chromosomes, analyzed at 1 Mb resolution, do not show TADs or compartments, they could not be related to the observed globular structures.

Different levels of chromatin organization have been observed, depending on the species analyzed and the methods used (reviewed in Kusnetzova and Sheval 2016). Several studies showed that in vivo chromatin is basically organized in 10-nm fibers and not 30-nm fibers, as found after fixation (reviewed in Maeshima et al. 2019). We measured a mean lower-order chromatin fiber diameter of ∼80 nm in DAPI-labeled barley metaphase chromosomes while scanning electron microscopy revealed 15- to 150-nm diameter fibers (Wanner et al. 1991), helium ion microscopy (with improved chromosome preparations preventing artifacts) revealed 11.6 -nm fibers (Sartsanga et al. 2021), and reticulate chromatin fibers with a diameter of up to 30 nm were observed in human metaphase chromosomes (Wako et al. 2020). Due to the visualization of fluorescent signals via a point spread function, signal thickness measurements (such as DAPI signals) might be overestimated, depending on the settings. Consequently, electron microscopy-based measurements are regarded as more reliable.

Within the chromatin fiber network, we found minor chromatin-free regions of ∼120×220 nm, but never larger cavities, as previously observed by electron microscopy in plant and human chromosomes (Hao et al. 1988, 1990; Hamano et al. 2014; Kuznetsova et al. 2017). Hamano et al. (2014) noted that the cavities were not visible when an ionic liquid method that keeps the chromosomes wet was used. Chen et al. (2017) also did not identify large cavities in human chromosomes. Thus, minor chromatin-free spaces occur within the condensed chromatin of mitotic chromosomes, whereas the previously observed larger cavities might be preparation artifacts (Beseda et al. 2020).

In conclusion, sister chromatids of large metaphase chromosomes in eukaryotes are formed by a helically arranged chromonema consisting of looped minor chromatin fibers, which may form clusters with small chromatin-free regions between them.

## Material and Methods

### Plant material

Barley (*Hordeum vulgare* L.) cv. Morex seeds were obtained from the Gene Bank of the Leibniz Institute of Plant Genetics and Crop Plant Research Gatersleben, Germany.

### Preparation and sequencing of chromosome Hi-C libraries

Suspensions of barley metaphase chromosomes were prepared from root tip meristems after cell cycle synchronization as described in Lysák et al. (1999) with the following modifications. Barley seeds were germinated for two days at 4°C, followed by incubation at 25°C for three days. The roots were fixed in 2% formaldehyde in 1×PBS buffer for 12 min at 5°C. Fixed metaphase chromosomes were released by mechanical homogenization into LB01 buffer (^79^Doležel et al., 1989) and stained by 4’,6-diamidino-2-phenylindole (DAPI) at a final concentration of 2 µg/ml. The chromosomes were purified by sorting using a FACSAria SORP flow cytometer (Becton Dickinson, San Jose, CA USA). The initial gating was performed on FSC-A vs. DAPI-A parameters. To discriminate doublets, the chromosome gate was drawn on a DAPI-A vs. DAPI-W scatterplot (Figure S16).

Three replicates of chromosome Hi-C libraries were prepared. For each replicate, five million chromosomes were flow-sorted into a 15-ml Falcon tube with 2 ml LB01, centrifuged at 500 g for 30 min at 4°C, and the supernatant removed except for 20 µl. The pelleted chromosomes were gently resuspended in the remaining supernatant, diluted with 8 ml ddH_2_0, and spun down at 500 g for 30 min at 4°C. The supernatant was removed except for 20 µl. The pelleted chromosomes were gently resuspended and the entire sample was transferred to a 0.2 ml Eppendorf tube. Further steps were carried out with an Arima Genomics Hi-C kit (Arima Genomics, San Diego, CA USA) according to the manufacturer’s protocol. The sequencing libraries were prepared with a NEBNext Ultra II DNA library preparation kit (NEB, Ipswitch, MA, USA) with 10 cycles of PCR amplification. Libraries were sequenced on an Illumina NovaSeq 6000 instrument (Illumina, San Diego, USA) in 150-bp paired-end mode. The parameters of the generated Hi-C data are summarized in Table S4.

The data for chromosome model simulation were generated by an alternative protocol. Eight million chromosomes were sorted into a 15-ml Falcon tube with 2 ml isolation buffer (Šimková et al. 2003), centrifuged at 500 g for 30 min at 4°C, and the supernatant removed except for 45 µl. Pelleted chromosomes were gently resuspended in the remaining supernatant and mixed with low-gelling agarose to create a 90 µl 1% agarose plug. The plug was washed twice for 30 min in 2 ml TE buffer (10 mM Tris-HCl, 1 mM EDTA, pH 8) on ice. Subsequent Hi-C library preparation steps, including DNA digestion, biotin fill-in, ligation, and chromatin release from the plug, were performed according to the “In situ Hi-C in agar plugs protocol” of Rao et al. (2014) with minor modifications: the DNA was digested by 800 U HindIII, and biotin-14-dATP was replaced by biotin-14-dCTP. The completed Hi-C library was sequenced on the HiSeq2500 system in 100-bp paired-end mode.

Hi-C reads were mapped against the barley cv. Morex genome version 2 (v2; Monat et al. 2019) using a previously described computational pipeline (Beier et al. 2017; Monat et al. 2019). Contact matrices were visualized in the R statistical environment (R Core Team 2020).

### Contact probabilities from Hi-C data

All contacting pairs from Hi-C data were counted and mapped to the Morex genome version 2 (v2), as a function of the genomic distance between them (*d*). All values of distance *d* were split into bins on a logarithmic scale from 100 kb to 1 Gb. The contact probability of each bin was calculated as the sum of all observed pairs within the distance range of this bin, divided by the number of all genomic loci within the same distance range.

### Quantification of Hi-C local contacts in metaphase chromosomes

The chromosomes were divided into 5-Mb-long non-overlapping regions. For each region, we considered only Hi-C pairs with at least one side of the contacting pair inside of this region or with both sides of this pair spanning this region (Figure S3). For each region, the pairs were counted based on the genomic distance between them (*d*). The counted pairs as a function of *d* are well described by the sum of an exponential and a Gaussian distribution. The Gaussian distribution coincides with the bump characteristic of a helical arrangement (Figure 1B-D), and its center indicates the turn length of the helical arrangement around the analyzed region. The center of the Gaussian distribution was assigned as the turn length for each region and was used to build the graphics in Figure 1E.

### Building the helical path with a changing radius and estimating the turn number

The monomers selected as the loop basis were constrained to a helical path in the Cartesian space. A turn length (in base pairs) *I_i_* was assigned to each monomer i according to the values calculated from the Hi-C data. With this value, we calculated how many monomers (*n_i_*) from the helical path would fit in this turn, with monomer size (in base pairs) *m* and the average major loop size (in monomers) *I_o_* (equation 1). Each monomer was then assigned a height *z_i_* and an angle *θ_i_* relative to the previous monomer. The height *z_i_* is equal to the constant turn height (400 nm) divided by *n_i_*, and the angle *θ_i_* is the full turn (360°) divided by *n_i_* (equation 2). The radius *r_i_* was calculated from the turn length (*n_i_* multiplied by the distance *d* between monomers) and the turn height as in equation 3, from the Pythagorean theorem. The cartesian coordinates were then retrieved by these cylindrical coordinates.

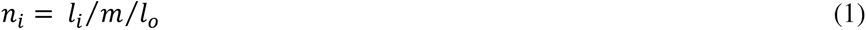

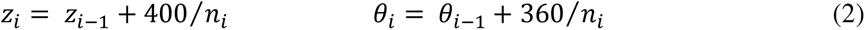

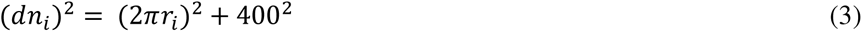

The number of turns is the number of cycles in the function *cos(θ)*, using 1 Mb as the monomer size and 1 monomer as the average major loop size.

### Plant growth and treatment for FISH

Barley seeds were incubated for three days in the dark at room temperature. Roots were collected in cold tap water and incubated on ice for 24 hours (h). The roots were transferred into an ethanol:acetic acid (3:1) fixation solution and placed under a vacuum for 5 min. The roots were stored in this solution overnight at room temperature (RT).

### Plant growth and treatment for EdU labeling of SCEs

Barley seeds were incubated for three days in the dark, followed by incubation in 20 µM EdU in Hoagland medium for 17 h at RT. The seedlings were transferred into fresh Hoagland medium and incubated for 19 h in the dark at RT (recovery phase). Roots were cut from the seedlings and incubated in ice-cold water for an additional 24 h. Finally, the roots were fixed in ethanol:acetic acid (3:1) for 5 min under a vacuum and overnight at RT.

### Chromosome spread preparation

Selected roots were washed twice in ice water for 5 min and once in a citric buffer for 5 min. The roots were digested in an enzyme mixture [1% Pectolyase (Sigma); 1% Cytohelicase (Sigma); 0.7% Cellulase R-10 (Duchefa), and 0.7% Cellulase (Calbiochem) in 0.1 M citric buffer] at 37°C for 45 min. Following enzymatic digestion, the roots were washed twice in ice-cold water for 5 min and twice in ice-cold ethanol for 5 min. Root tips without root caps were collected into the tube containing ethanol:acetic acid (3:1) and mashed. 7 µl of this suspension was dropped onto a cold wet slide, transferred to a hot plate (55°C), and fixed with 25 µl of ethanol:acetic acid (3:1). The slides were allowed to dry at least for 1 h at RT.

To preserve the native volume of the chromosomes, metaphase chromosomes were flow-sorted into 1× meiocyte buffer A (1× buffer A salts, 0.32 M sorbitol, 1× DTT, 1× polyamines) and subsequently embedded into a 5% polyacrylamide gel as described (Bass et al. 1997; Howe et al. 2013) with minor modifications (Němečková et al. 2020).

### FISH

To detect half and full helix turns within chromosome arm 5HL by fluorescence microscopy, oligo-FISH probes were designed against the chromosome-scale sequence assembly of the barley cv. Morex genome assembly version 1 (Morex v1; Beier et al. 2017; Mascher et al. 2017) using Abor Biosciences’ proprietary software. Briefly, target sequences were cut into 43–47 nucleotide-long overlapping probe candidate sequences that were compared to the rest of the genome sequence to exclude any candidates with potential cross-hybridization based on a predicted *Tm* of hybridization. Non-overlapping target-specific oligonucleotides were selected for the final sets and synthesized as myTAGs® Labeled Libraries (Arbor Bioscience, Ann Arbor, MI USA).

To validate the lengths of intervals covered by the particular oligo-FISH probes, we verified the completeness of the Morex v2 genome assembly in the analyzed region of 5H using an optical map constructed from barley cv. Morex on the Saphyr platform of Bionano Genomics (Figure S8). The optical map contigs were aligned to the selected interval (442–599 Mb) in the chromosome 5H pseudomolecule using RefAligner version 9232 (Bionano Genomics). Query-to-anchor comparison was performed with default parameters and a p-value threshold of 1e−10. The alignment was visualized in Bionano Access version 1.5. The coordinates from the original design were replaced and are now based on the improved Morex assembly v2 (Monat et al., 2019). The labeled region starts at position 442.1 Mb. For simplicity, the designed probes were given the names of different bird species and labeled in different colors: Stork-Atto647N, Eagle-Alexa488, Ostrich-Atto594, Rhea-Alexa488, Moa-Atto594, and Flamingo-Alexa488 (Figure 2C; Figures S6, S7; Table S2).

Subtelomeres were labeled with the HvT01 oligo-probe (Belostotsky and Ananiev 1990; Schubert et al. 1998) using TexasRed, and telomeres were labeled with the *Arabidopsis* telomere-type oligo probe (Richards and Ausubel 1988) using Cy5. The centromeres were labeled with the probe for centromeric repeats (GA)15 (Zhang et al. 2019) using FAM, and the NORs were labeled with the 45S rDNA probe using the clone pTa71 and Alexa488 (Gerlach and Bedbrook 1979).

For FISH, the slides were incubated in 45% acetic acid at RT for 10 min and washed in 2×SSC at RT for 10 min. When strong background signals due to cytoplasm were present, 50 µl of pepsin solution (0.1% in 0.01M HCl) was added to the slide, which was then covered with parafilm and incubated in a wet chamber at 37°C for 10 min. The slides were washed twice in 2×SSC for 5 min and post-fixed with 4% formaldehyde in 2×SSC at RT for 10 min. The slides were then washed 3 times in 2×SSC at RT for 4 min and immediately dehydrated in an ethanol gradient (70%, 85% and 100%, 2 min each). The slides were air-dried for at least one hour. In the meantime, all selected oligo probes were pooled into a microtube and the solutions evaporated using a SpeedVac concentrator (Eppendorf). The probes were reconstituted in 1 µl of ddH_2_O and 19 µl of hybridization mixture (50% formamide; 10% dextran sulfate; 10% salmon sperm DNA; 2×SSC). The entire volume of the reconstituted probe was added to the slide, covered with a coverslip, and sealed with rubber cement. The slides were incubated for 20 min at 37°C and denatured on a hot plate (70°C) for 3 min. Finally, the slides were directly placed into a wet chamber and hybridized for 22 h at 37°C. Post-hybridization washing was carried out as follows: the slides were briefly washed in 2×SSC at RT to remove the coverslip, washed with shaking at 58°C for 20 min, followed by 5 min in 2×SSC at RT, dehydrated in an ethanol series (70%, 90% and 96%, 2 min each), air-dried in the dark, and counterstained with 8 µl DAPI (2 µg/ml in Vectashield).

### Combination of EdU labeling and FISH

Seedling growth and treatment were performed as described for the EdU labeling experiment. Slides containing metaphase spreads with EdU-labeled SCEs were pre-selected and used for oligo-FISH. Depending on the cytoplasm background, the slides were incubated in 45% acetic acid at RT for 10 min and washed in 2×SSC at RT for 10 min or directly post-fixed with 4% formaldehyde in 2×SSC at RT for 10 min. The subsequent steps were identical to the FISH procedure described above.

### Super-resolution microscopy and measurement of FISH signal volume

To detect the ultrastructural chromatin organization of chromosomes at a resolution of ∼120 nm (super-resolution achieved with a 488 nm laser excitation), spatial structured illumination microscopy (3D-SIM) was performed with an Elyra PS.1 microscope system with a 63×/1.4 Oil Plan-Apochromat objective using ZENBlack software (Carl Zeiss GmbH). Images were captured separately for each fluorochrome using the 642, 561, 488, and 405 nm laser lines for excitation and appropriate emission filters (Weisshart et al. 2016). Maximum intensity projections of whole cells were calculated using ZENBlack software. Zoomed-in sections were presented as single slices to indicate the subnuclear chromatin structures at super-resolution. 3D rendering to produce spatial animations was performed based on SIM image stacks using Imaris 9.6 (Bitplane) software. The FISH signal and DAPI-labeled whole chromosome volumes were generated and measured with the Imaris tool ‘Surface’.

### Polymer simulation

A simulation for barley metaphase chromatin organization was performed based on a 10 nm chromatin fiber and the bottle-brush model of chicken containing a protein scaffold as described in Gibcus et al. (2018). We assumed that the individual 400 nm fiber turns consisted of larger and smaller flexible loops. Therefore, a partial intermingling of the minor loops originating from adjacent turns occurred (Figure S10).

In all simulations, the chromatin was modeled as a beads-on-a-string homo-polymer. Molecular Dynamics simulations (MD) were performed using the OpenMM Python application programming interface (Eastman et al. 2017) and the OpenMM-lib library (https://github.com/mirnylab/). The motion of the polymer was simulated based on Langevin dynamics, with a temperature of 300 K, a collision frequency of 0.001 ps^-1^, and a variable time step (Gibcus et al. 2018). In this simulation, three internal forces are applied to the polymer: (1) a harmonic force covalently binding two neighboring beads; (2) a harmonic angular force between three sequential beads with a spring constant of l k_B_T/rad^2^; and (3) a polynomial repulsive force allowing for the crossing of the fiber by setting an energy truncation value of 1.5 k_B_T, when the distance between two non-bonded beads is zero. In the bottle-brush model, the entire polymer is organized into side-by-side major loops divided into side-by-side minor loops, as proposed by Gibcus et al. (2018). The lengths of both types of loops are randomly chosen following an exponential distribution. Two external forces are applied to constrain the polymer in a cylindrical bottle-brush model: (1) the monomers forming the base of the major loops are tethered to a helical path by a harmonic potential with a spring constant of 4 k_B_T/nm^2^; and (2) this helical path is at the center of a cylinder, whose boundaries are defined by a harmonically increasing potential with a spring constant of 10 k_B_T/nm^2^. Apart from these characteristics, the bottle-brush model is defined by four parameters: (1) the average length of major loops; (2) the average length of minor loops; (3) the turn height, and (4) the turn length of the helical path. These parameters are explicitly mentioned for each model we tested (Figure S12C). All simulations start with a conformation where the base of the major loops follow a calculated helical path, and the loops emerge radially. All monomers in the helical path are assigned cylindrical coordinates (angle, radius, and height). These coordinates are calculated so that the monomers are equally separated given the specified total number of major loops and the turn height and converted to Cartesian coordinates. The simulation runs until the contact probability is equilibrated.

To reproduce the contact probability of chicken chromosome 1 at prometaphase (30 min after the release of G2 arrest), as described by Gibcus et al. (2018), with major loops of 400 kb (average length), minor loops of 80 kb (average length), 150 nm turn height, and 8 Mb turn length, we used a coarse-grained model where each monomer represents one nucleosome. Covalent bonds were separated by 10 nm average distance and 1 nm wiggle distance, cylindrical confinement secured a volume of 11^3^ nm^3^ per monomer, and the monomers in the helical path were separated by 50 nm. Using the same parameters, we built four models to reproduce the contact probability of metaphase barley chromosome 5H, varying the average length of major and minor loops, the turn height, and the turn length. These models simulate a region with 5 turns of the helical path (Figure S12D, E). To compare helical and half-helical arrangements, we used the same coarse-grained model described above but simulated a 10-Mb region with 2-Mb turn length, 100-nm turn height, 100kb major loops, and 10-kb minor loops. The radius of the helical path was set to 100 nm, and no cylindrical confinement was set. The half helical arrangement differed from the helical arrangement only in terms of the position of the major loops’ basis: only the absolute values of the x-coordinate were considered in the conversion from cylindrical to Cartesian coordinates. Ten replicates per model were used to calculate the contact probability.

To observe the changing turn length, we simulated the ∼600 Mb of chromosome 5H with a coarse-grained model where each monomer represents 2 kb. In agreement with the volume measured for the entire chromosome (∼43^3^ nm^3^/2 kb), we set covalent bonds that were separated by an average distance of 40 nm and a wiggle distance of 4 nm. The cylindrical confinement was set separately for each major loop, i.e., 800 nm in height around the position of the loop basis and radius proportional to the turn length, with a minimum of 500 nm and a maximum of 900 nm following our chromatid width measurements after EdU labeling (Movie 12). Major and minor loops were 1 Mb and 400 kb long, respectively. Monomers in the helical path were separated by 50 nm. The helical path was calculated with a constant turn height of 350 nm and a radius proportional to the local turn length, as previously calculated from our Hi-C data. At the constriction regions, where no helical arrangement occurs, the position of the major loops’ basis follows a straight line on the z-axis and minor loops are only 40 kb long, a value that approximately mimics the observed size of the primary constriction.

### Contact probabilities of polymer models

To measure the contact probabilities of the equilibrated models, the final conformation of each simulation was calculated by counting all monomers spatially close to each other by no more than 51 nm, as in (^34^Gibcus et al., 2018). These pairs of monomers were then grouped according to their distance in the linear genome. The sum of observed contacts in each group was divided by the sum of all possible contacts between two monomers separated by this group distance in the linear genome. The contact probability as a function of the linear genomic distance was directly compared to the experimental data obtained by Hi-C experiments upon normalization of all probability functions to be equal to 1 for 100 kb.

## Acknowledgments

This work was supported by the Deutsche Forschungsgemeinschaft (Schu 762/11-1), DFG-GACR project (award No. 18-14450J) and by the ERDF project ‘‘Plants as a tool for sustainable global development” (No. CZ.02.1.01/0.0/0.0/16_019/0000827). We thank Katrin Kumke and Ines Walde for excellent technical assistance, and Ingo Schubert for critical reading of the manuscript.

---

**Figure S1.**
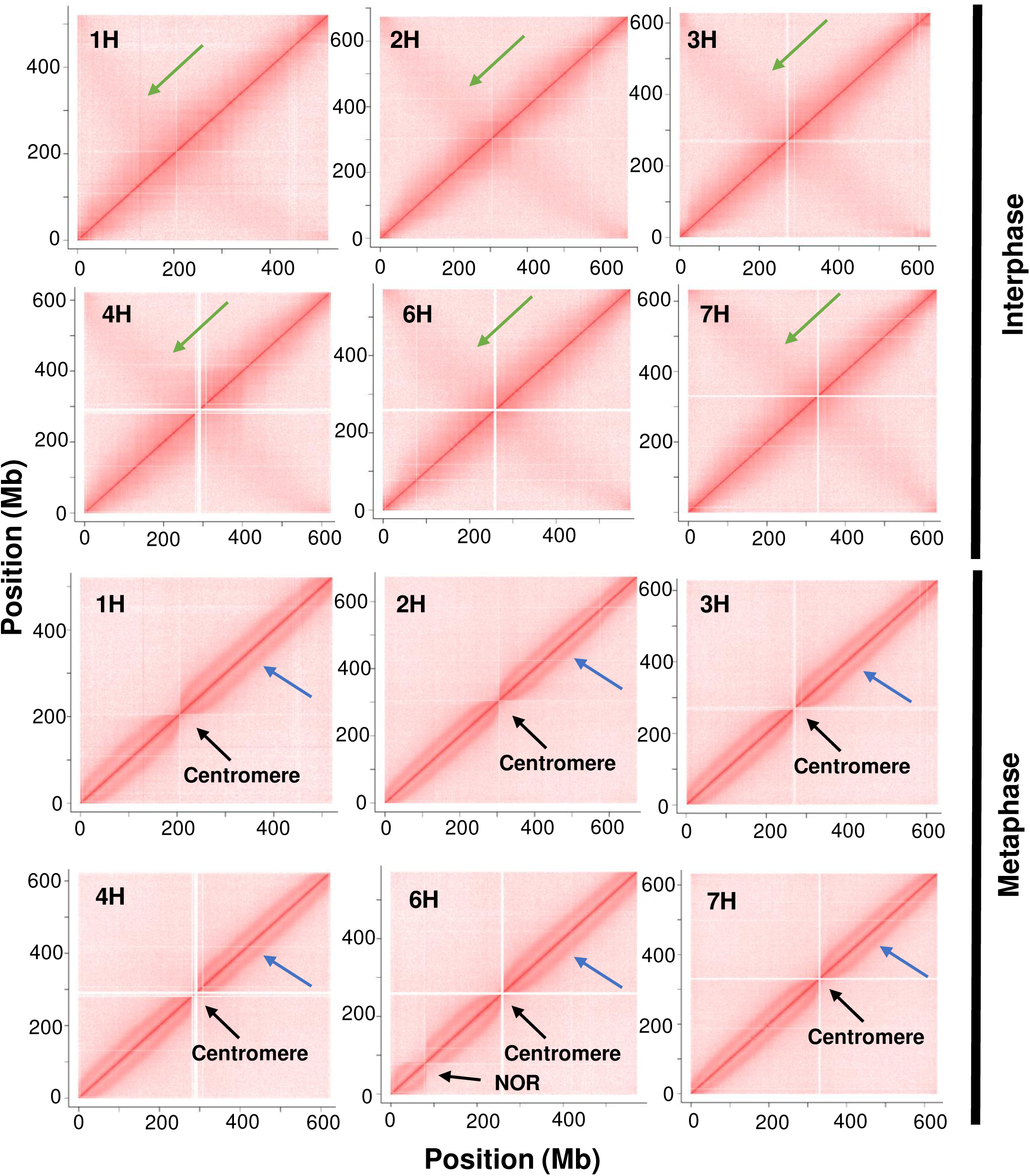
Hi-C contact matrices of all barley chromosomes (except 5H shown in Figure 1A) at interphase and metaphase from the first replicate of the Hi-C experiment. The anti-parallel diagonal in interphase (green arrows) indicates the Rabl configuration, while metaphase chromosomes show a parallel diagonal (blue arrows), indicating that a periodical contact pattern is missing at the NOR of 6H and the centromeres (black arrows).

**Figure S2.**
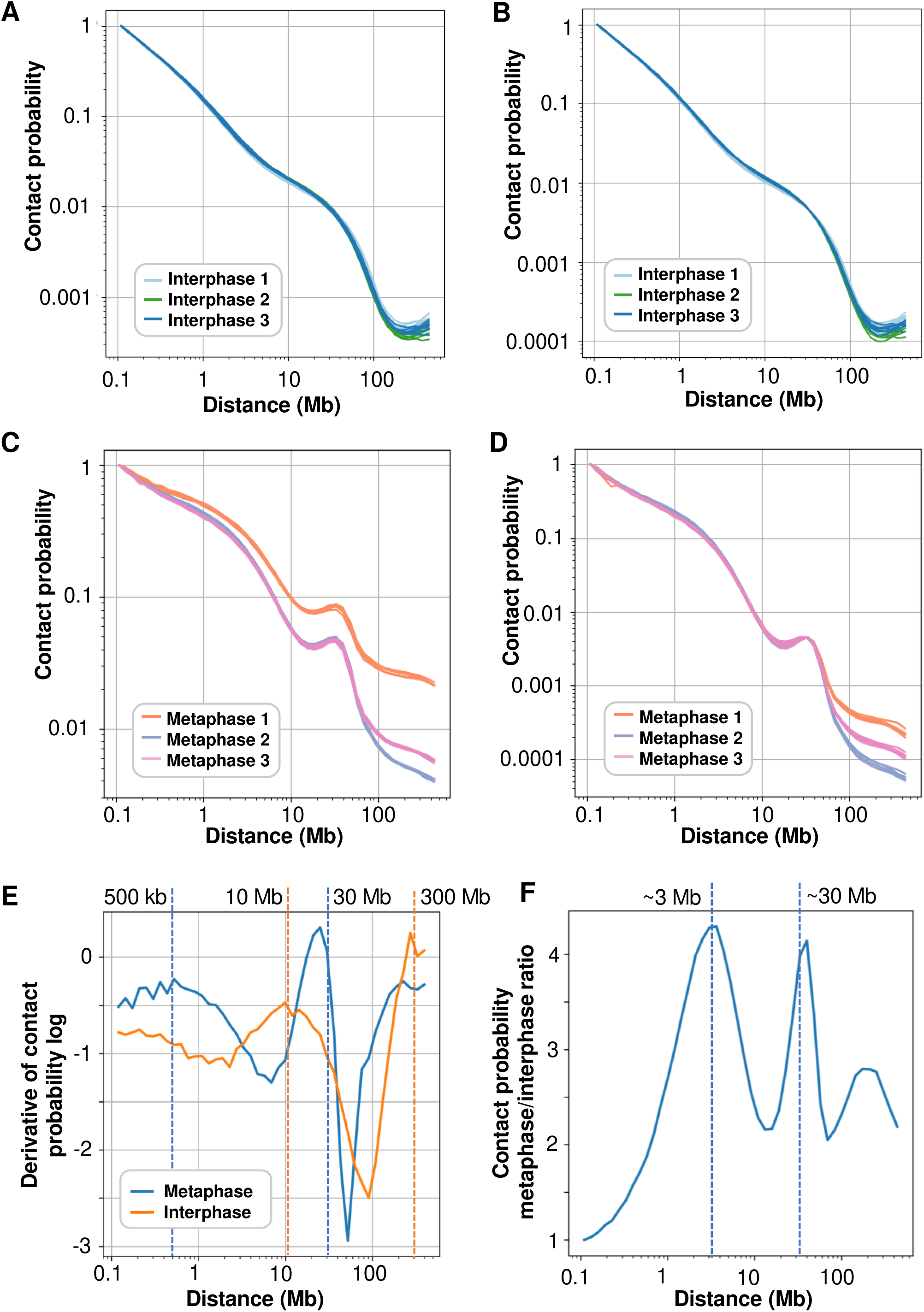
Contact probabilities calculated from the three Hi-C experimental replicates of interphase chromatin (A, B) and metaphase (C, D) chromosomes. In all graphics, each line represents one chromosome and each color one replicate. In **(A)** and **(C),** the probabilities are normalized to have the same value at 100 kb. In **(B)** and **(D),** the probabilities are exponentially normalized to have the same values at 100 kb and 30 Mb. In **(D),** the exponential normalization withdraws experimental variability and highlights the position of the bump characteristic of the metaphase contact probability. (**E**) Derivative of the contact probability log comparing metaphase **(D)** and interphase **(B).** (**F**) Contact probability ratio between metaphase **(D)** and interphase **(B)**. The dashed lines in **(E, F)** correspond to the observed peaks. In interphase, both peaks relate to the Rabl conformation reminiscent of anaphase. In metaphase, the first peak corresponds to the size of chromatin loops and the second to the size of helical turns. In the ratio of metaphase/interphase contact probabilities, the peaks highlight the differences between metaphase and interphase conformations.

**Figure S3.**
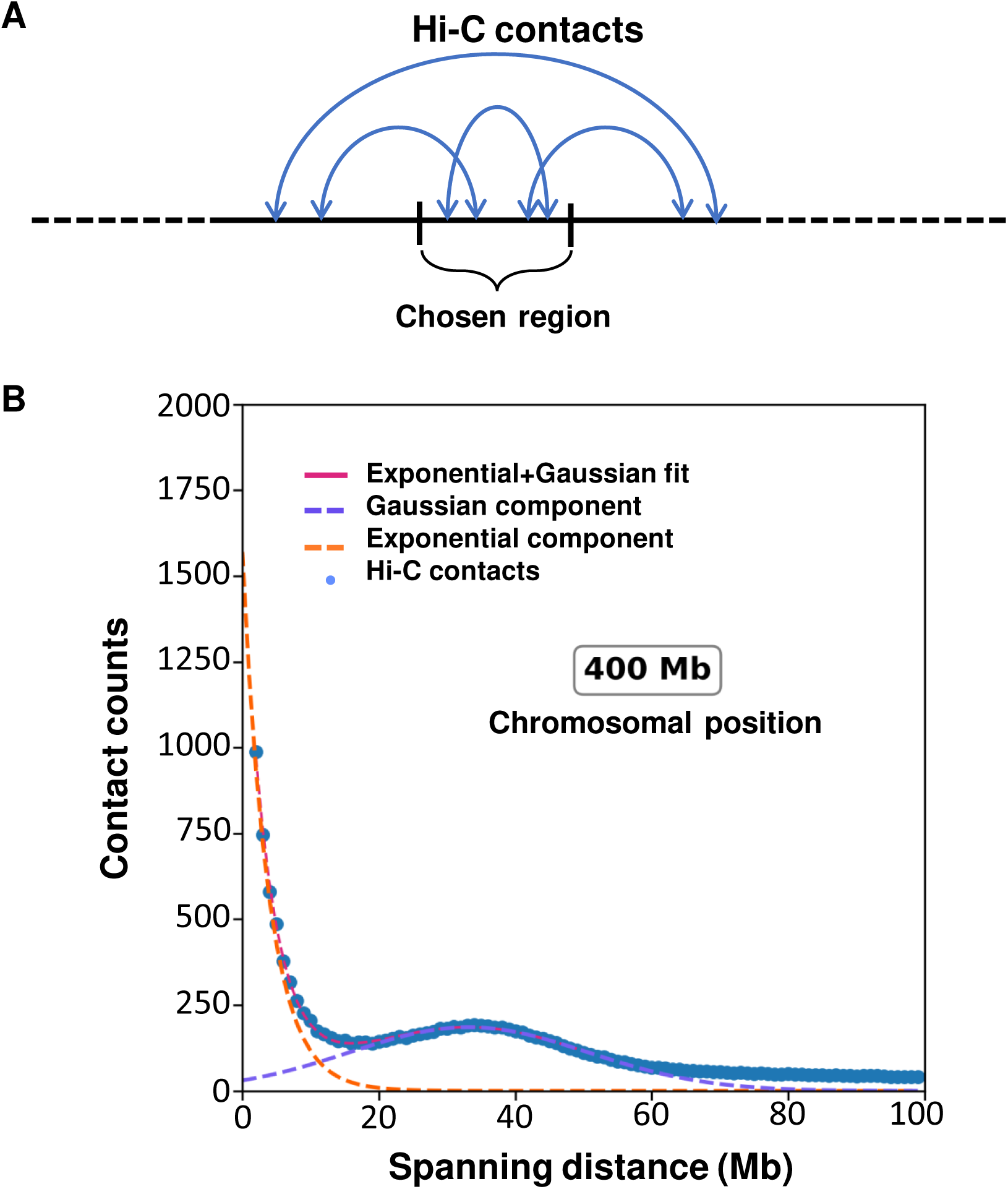
Quantification of Hi-C local contacts in metaphase chromosomes. The chromosomes were divided into 5-Mb-long non-overlapping regions. All Hi-C pairs within or spanning each region **(A)** were analyzed as a function of the distance between the contacting sites. For each region, this function was plotted as the number of contacts separated by 1 Mb, 2 Mb, … until 100 Mb **(B)**. Normalized, the results can be interpreted as a local contact probability (similar to Figure 1B). This function is well described by the sum of an exponential and a Gaussian distribution. The Gaussian distribution coincides with the bump characteristic of a helical arrangement (see Figure 1). Its center indicates the turn length of the helical arrangement around the analyzed region, which was used to build the graphics in Figure 1C.

**Figure S4.**
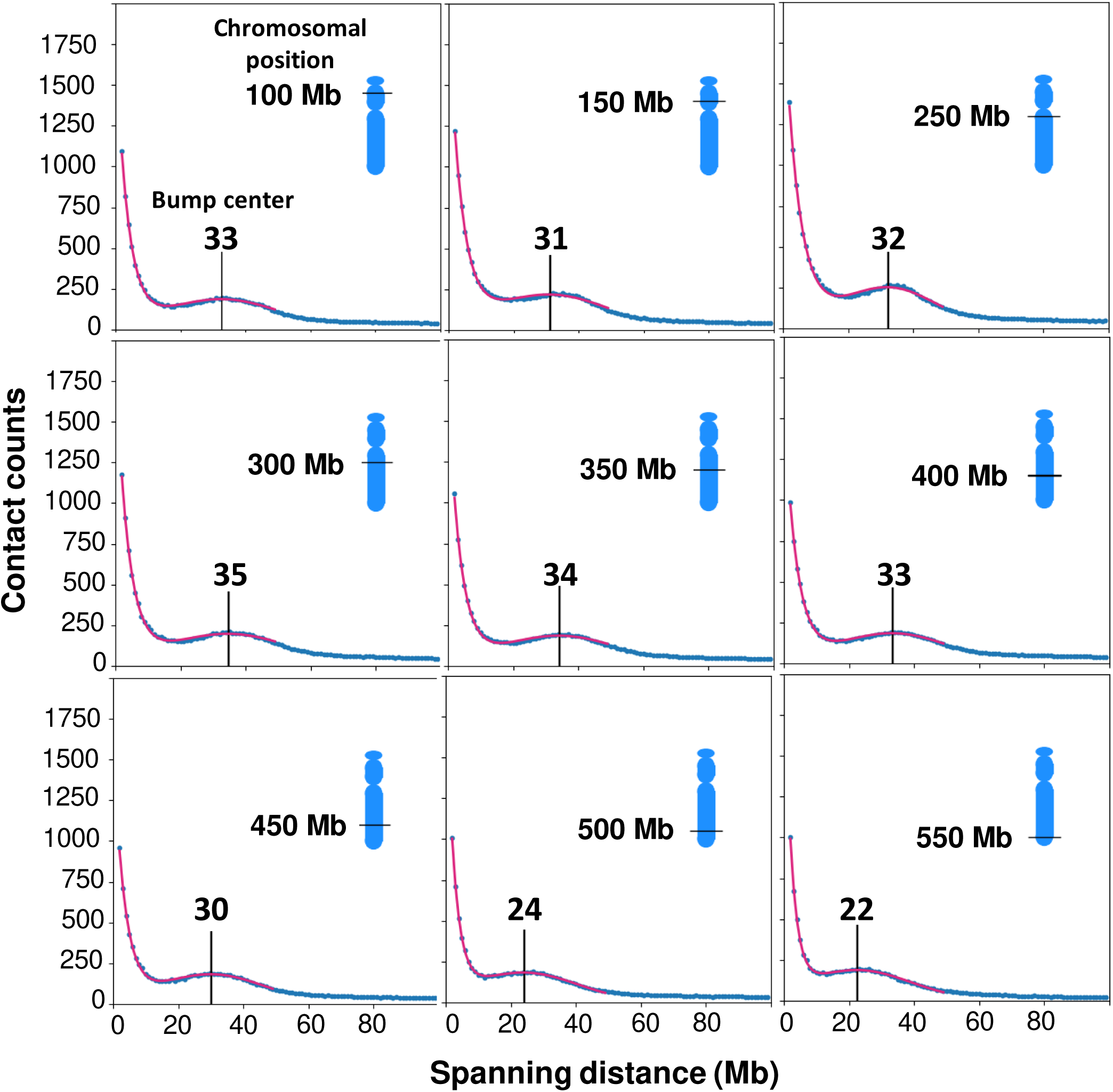
Number of local contacts at nine different positions (100–550 Mb) of chromosome 5H retrieved from experimental Hi-C data. At all positions, a prominent bump is present, with the center at slightly different positions. The bump represents a contact increase followed by a decrease at half of its position. This pattern indicates that the chromatin is arranged into turns, with turn length corresponding to the center of the bump. The bump position at any region along chromosome 5H was calculated as described in Figure S3. The placement of the bump for all 5 Mb regions (119), separated by 5 Mb along the 5H sequence, is visualized in Movie 1.

**Figure S5.**
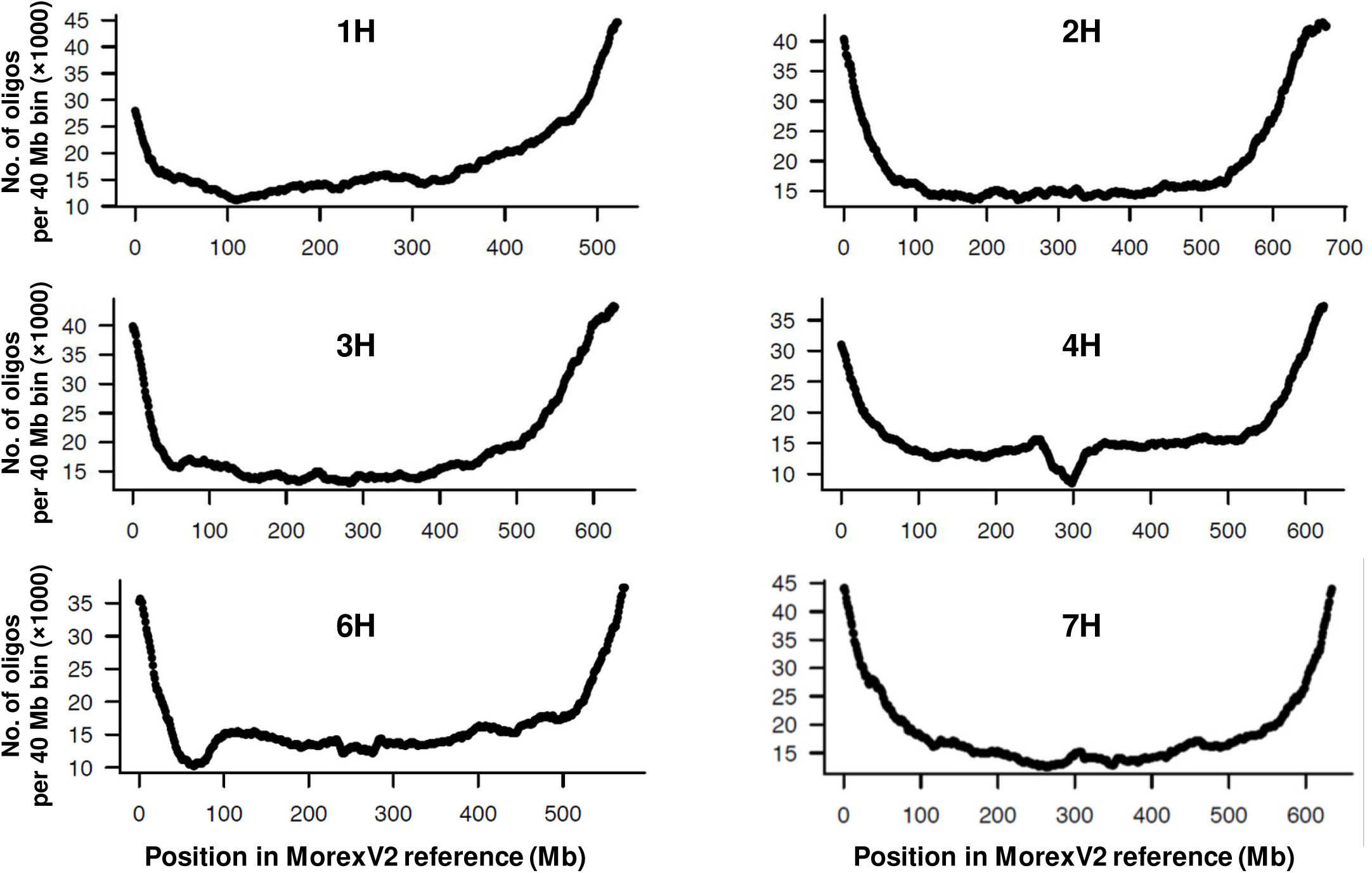
Predicted oligo density along all barley chromosomes (except 5H shown in. Figure 2A**) used to design the oligo-FISH experiment.** The x-axis shows the positions of the oligos on the chromosome in the Morex v2 genome. The y-axis represents the number of oligos within 40- Mb bins.

**Figure S6.**
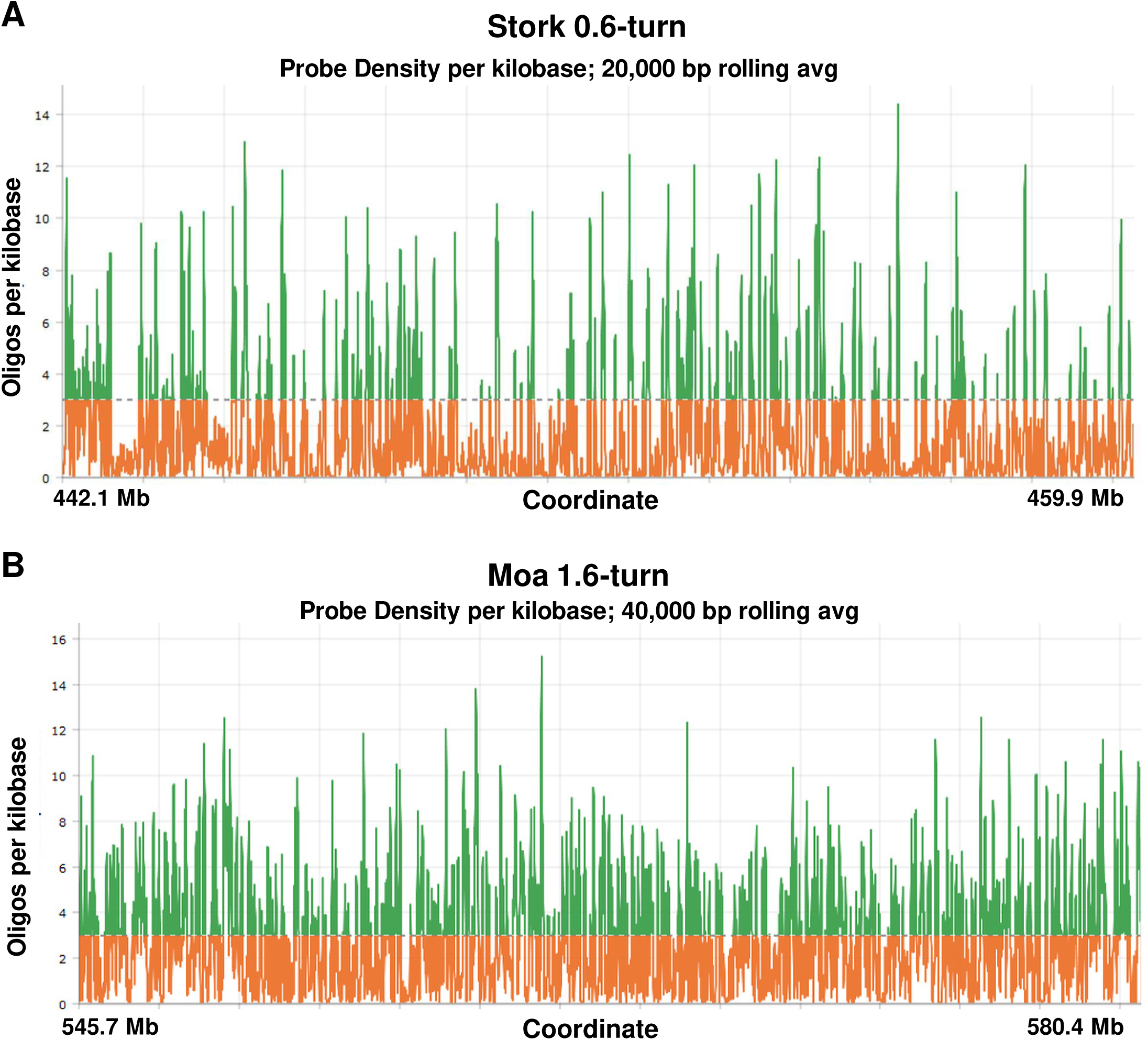
Graphical representation of the oligo density based on the barley cv. Morex v2 assembly covering the Stork and Moa regions subjected to oligo-FISH. **(A)** The 33,987 oligos of 45 nucleotides (on average) of the Stork probe homogeneously cover a 0.6-turn of the helix, with minor gaps arising from the lack of unique sequences. **(B)** The 108,831 oligos of 45 nucleotides (on average) of the Moa probe homogeneously cover a 1.6-turn of the helix without significant gaps. Orange marks the threshold of a minimum of three oligos per kilobase. Green denotes a sufficient number of oligos for FISH in the selected region.

**Figure S7.**
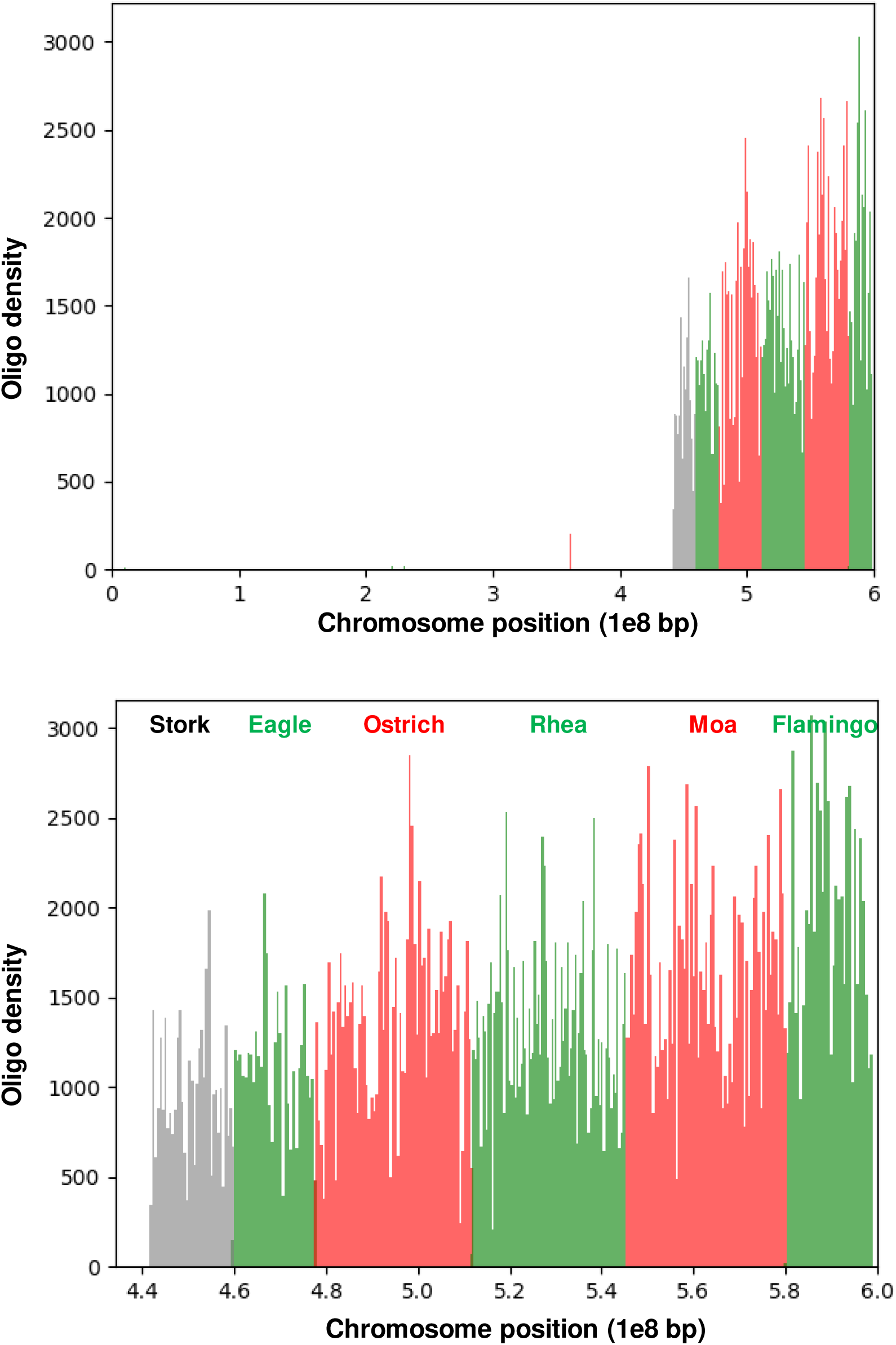
Graphical representation of the oligo densities of all designed probes at the long arm of chromosome 5H. The lower diagram shows an enlarged view of the region covered by the oligo probes.

**Figure S8.**
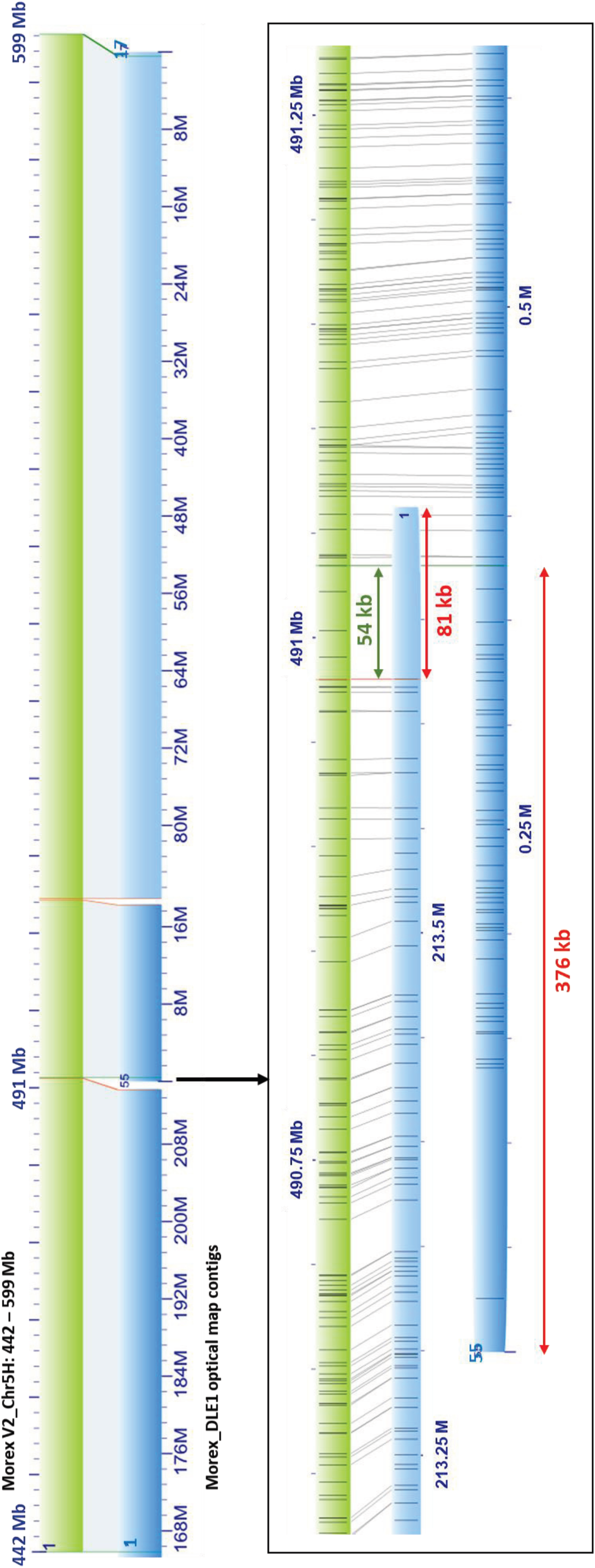
Validation of chromosome 5H interval 442–599 Mb in the Morex V2 genome assembly by optical mapping. The selected sequence segment (green bars) shows good alignment to the optical map contigs 1, 55, and 17 (blue bars), indicating the overall correctness and completeness of the sequence (top), except for a region around 491 Mb, where non-aligned overhangs of contigs 1 and 55 indicate a missing sequence of at least 403 kb (bottom).

**Figure S9.**
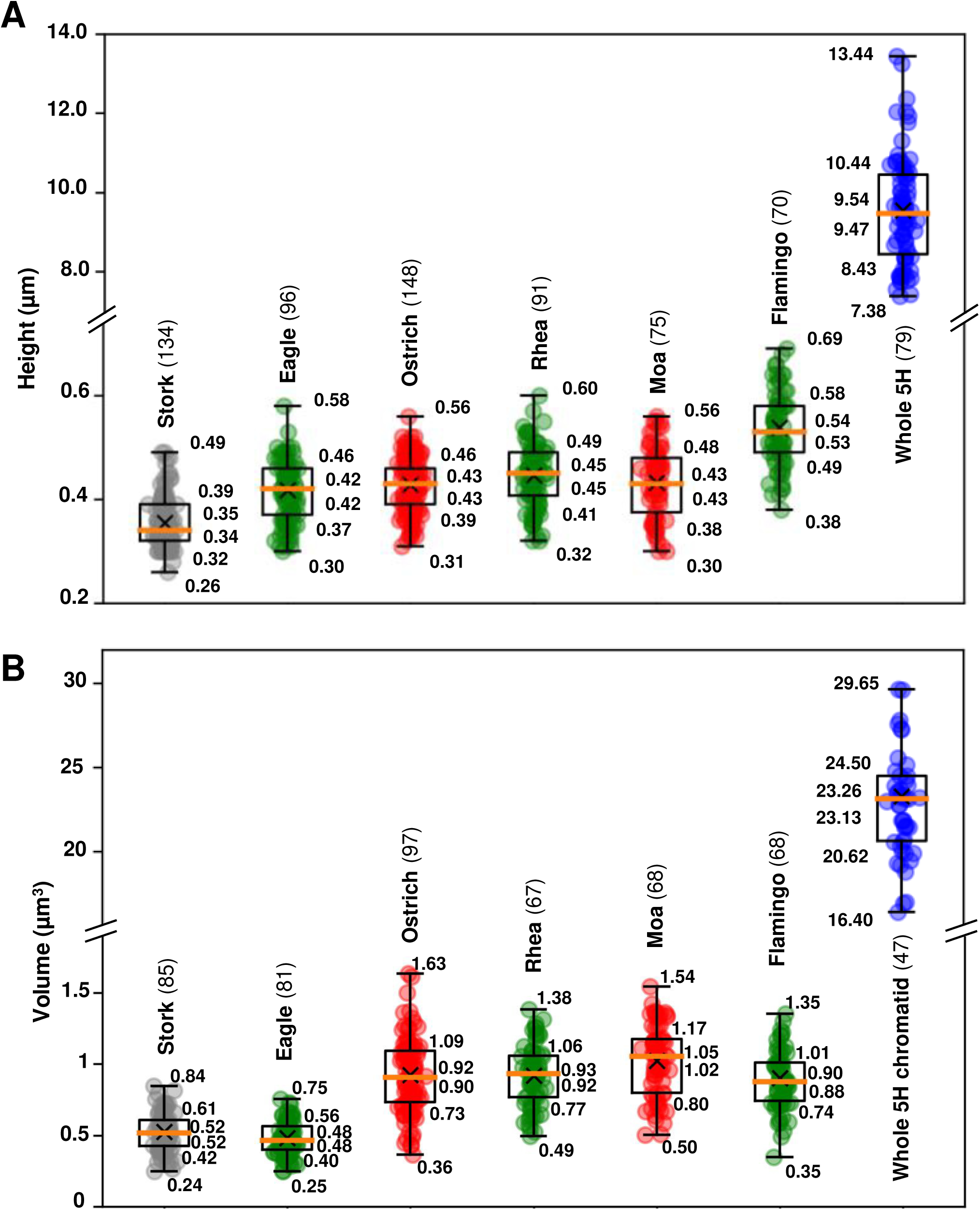
Box-plots showing the measured signal heights (A) and volumes (B) of the oligo-labeled regions on chromosome 5HL. The variable heights of the probes are consistent with the different turn lengths along the mitotic chromosome (Figure 2) and the different degrees of metaphase chromosome condensation. Volumes were measured after surface rendering using Imaris 9.6, and the values correspond to a single chromatid. The numbers of measured chromosomes per oligo probe are shown in parentheses.

**Figure S10.**
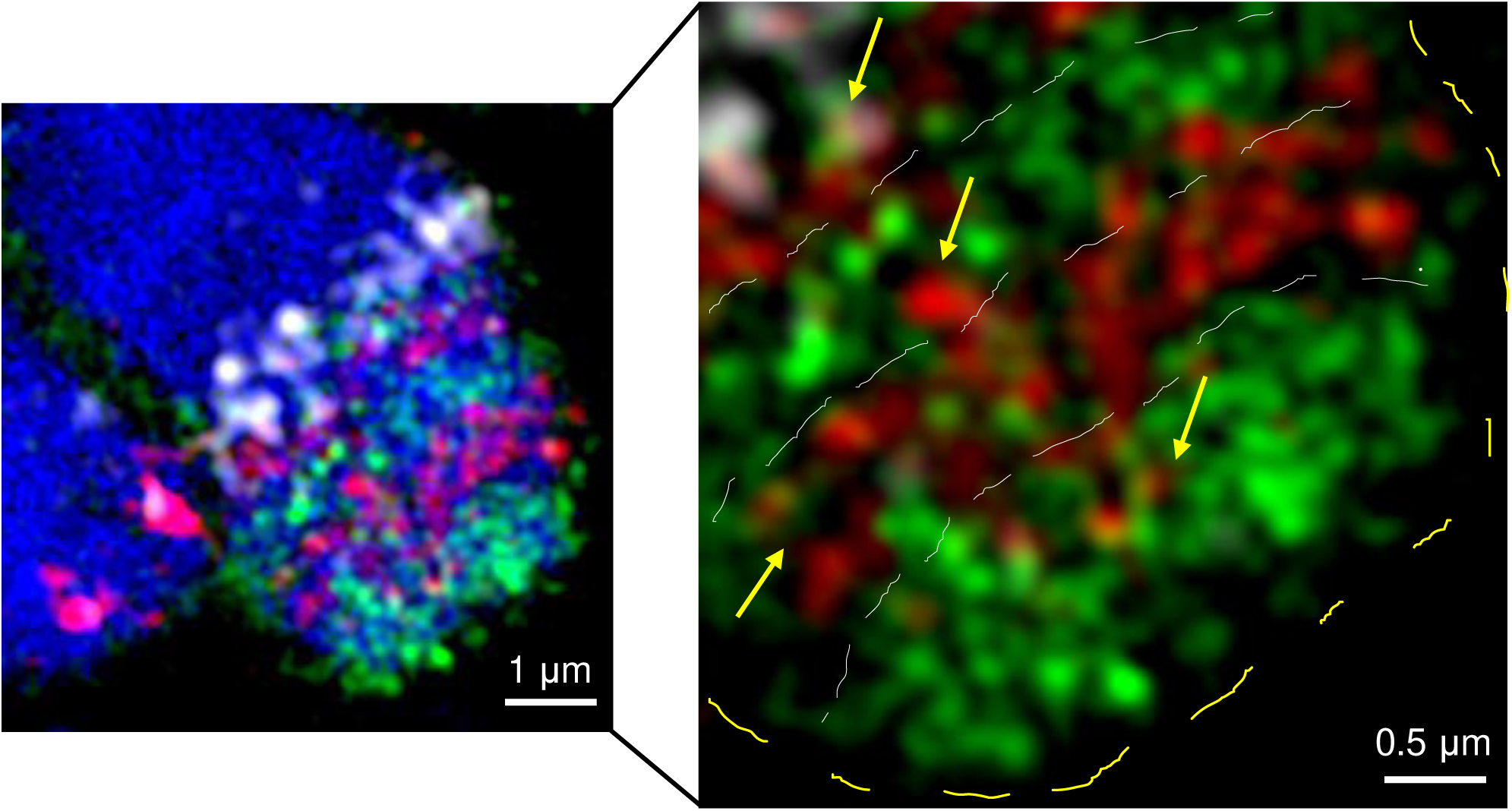
Adjacent helix-turns are partially intermingled. The differentially oligo-FISH labeled turns at chromosome arm 5HL intermingle. Smaller looped chromatin fibers forming the turned chromonema (indicated by dashed lines and marked by different colors) invade each other’s space (arrows in the right enlarged image). Chromatin was counterstained with DAPI (blue).

**Figure S11.**
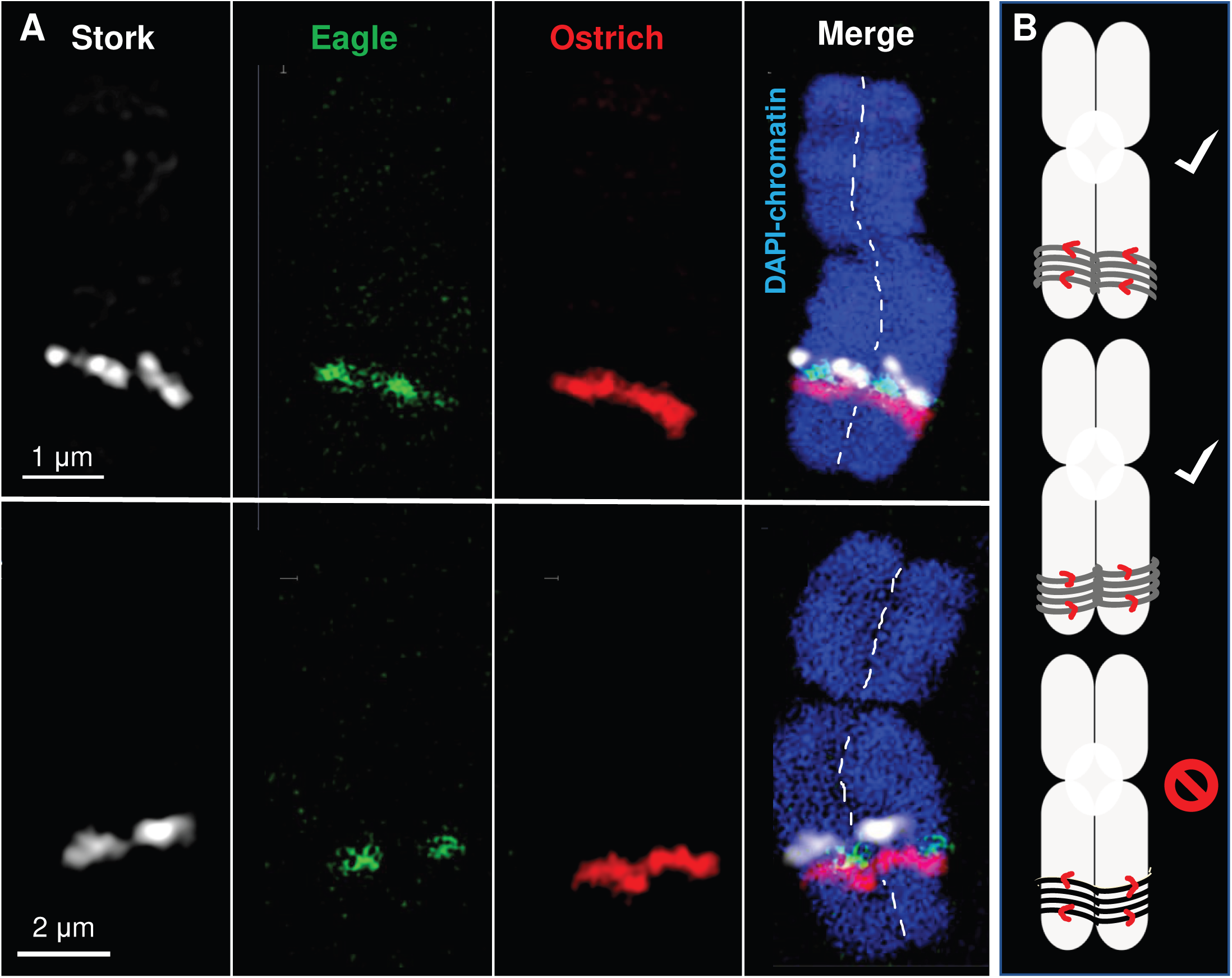
In both chromatids, the 400-nm fibers turn in the same direction. **(A)** The oligo-FISH probes (two examples of Stork, Eagle, and Ostrich are shown) reveal that the sister chromatids coil in the same direction, and not in a mirrored manner. In the merged image, the sister chromatids are divided by a dashed line. **(B)** The schemata show arrangements that were observed (top and middle) and not observed (bottom).

**Figure S12.**
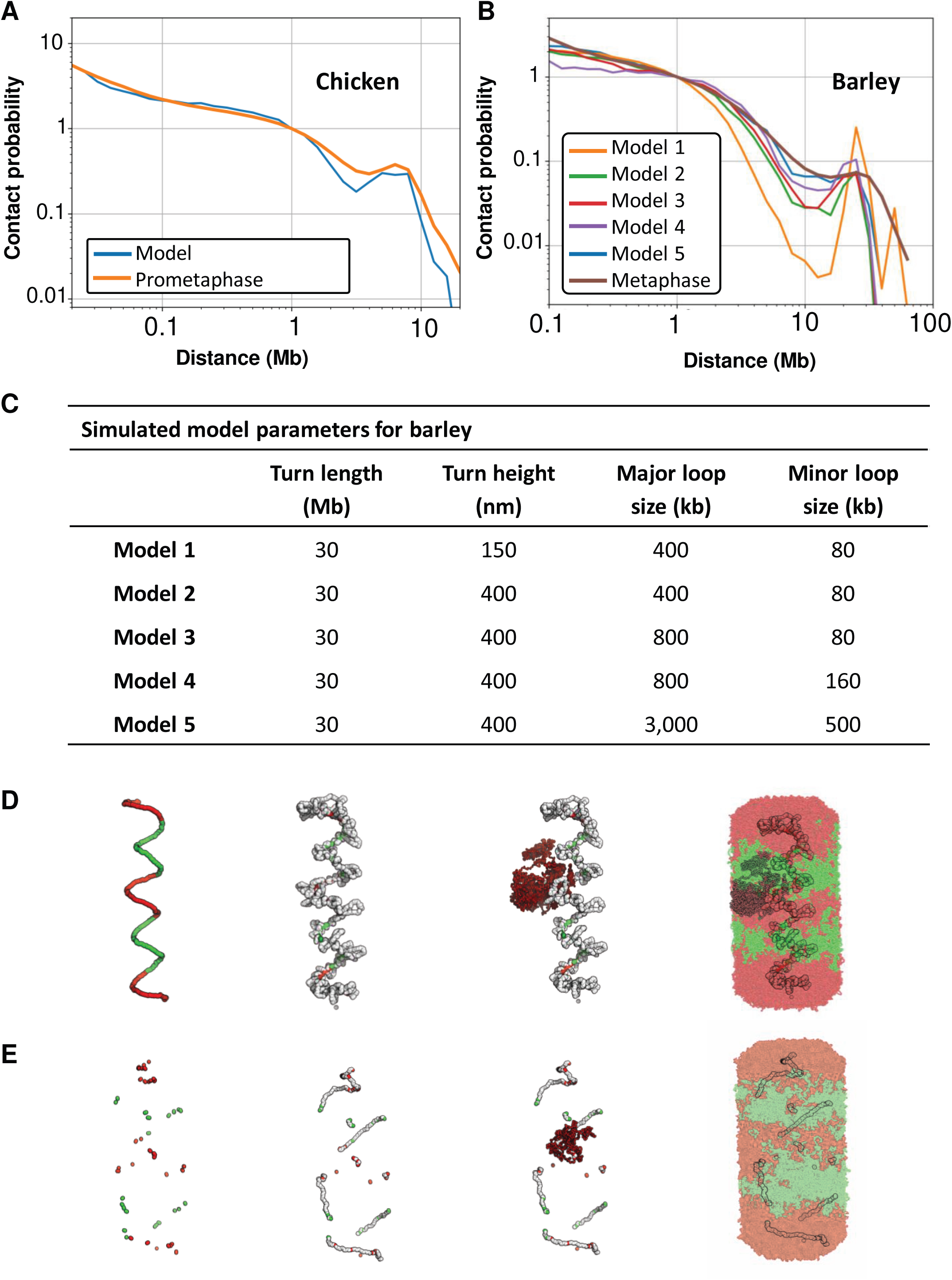
Comparison of applied bottle brush models for condensed barley vs. chicken chromosomes. **(A)** Contact probability of a barley polymer model generated with the settings described by Gibcus et al. (^34^2018) compared with the Hi-C contact probability for chromosome 1 of chicken at prometaphase (^34^Gibcus et al., 2018). **(B)** Contact probabilities of four barley polymer models, with the same settings as in **(A)**, but with varying helical arrangement parameters as shown in table **(C)**. The models are compared with the Hi-C contact probability of barley chromosome 5H at metaphase. All probabilities are normalized to be equal to 1 at 100 kb. **(D, E)** Conformations of barley models 3 **(D)** and 5 **(E)**. The 30-Mb turns are colored in alternating red and green. From left to right: the bases of major loops form the helical path; the bases of the minor loops are shown in white; a single major loop is divided into minor loops; all loops together complete the model. In **(D)** the bases of major loops (model 3) follow a helical path where the monomers are ∼10 nm apart. In **(E),** the bases of the major loops (model 5) follow the same path, but the monomers are ∼100 nm apart. Thus, the helical structure is not as constrained as in **(D)**. Consequently, in **(D)**, a backbone scaffold is formed. In **(E)**, the scaffold appears dispersed.

**Figure S13.**
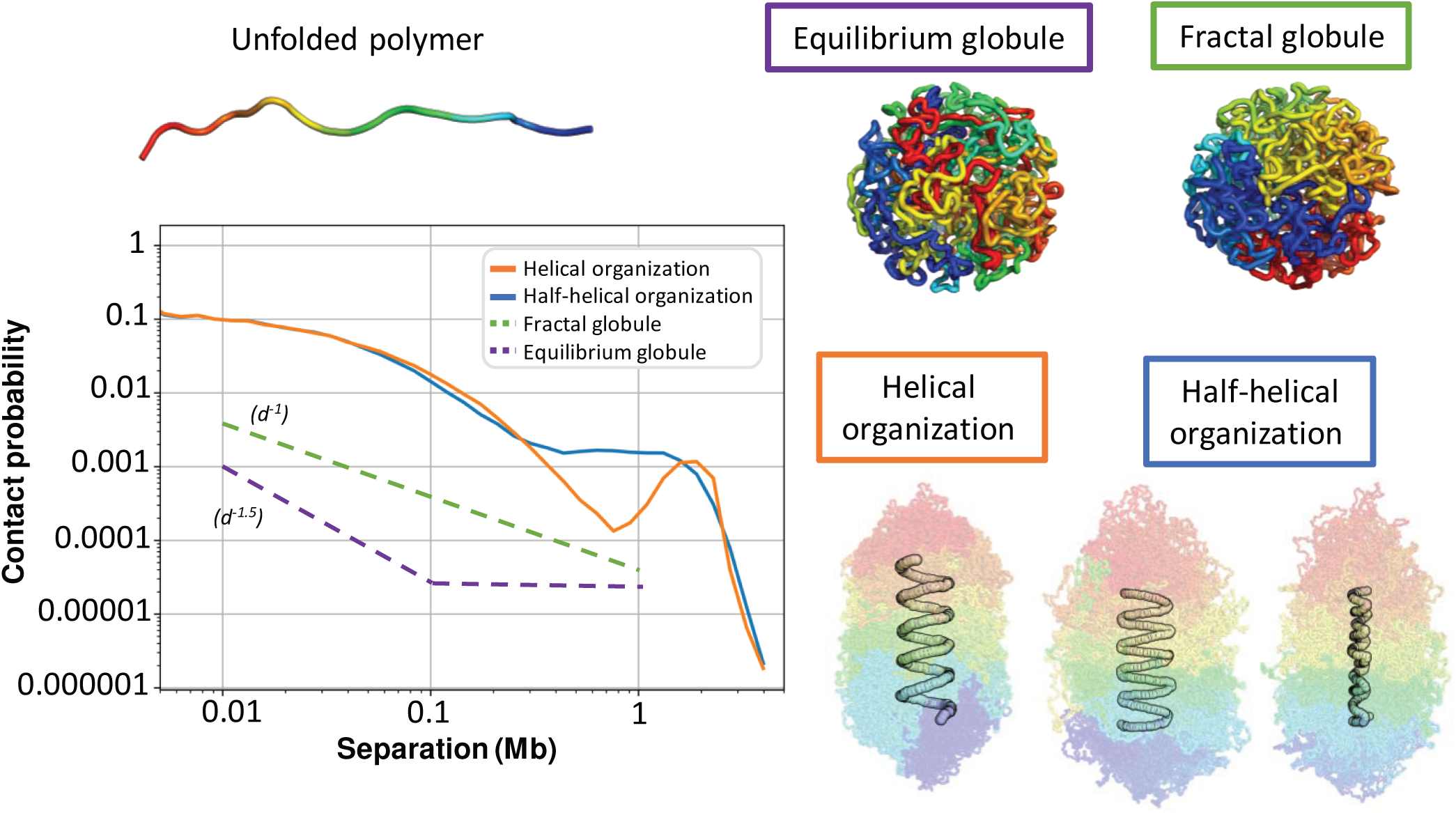
Contact probabilities of polymer models support a helical chromatid organization. The diagram shows the contact probabilities of different polymer models predicted for equilibrium and fractal globules, for simulated helical organization, as suggested by Gibcus et al. (2018), and for a simulated half-helical arrangement with changing handedness, as suggested by Chu et al. (2020). In all spatial models, the polymer is colored as shown for the unfolded polymer. For the helical and half-helical organizations, the polymer is transparent, and the basis of the loops appears as white spheres arranged in turns. Only the helical model creates the contact probability bump.

**Figure S14.**
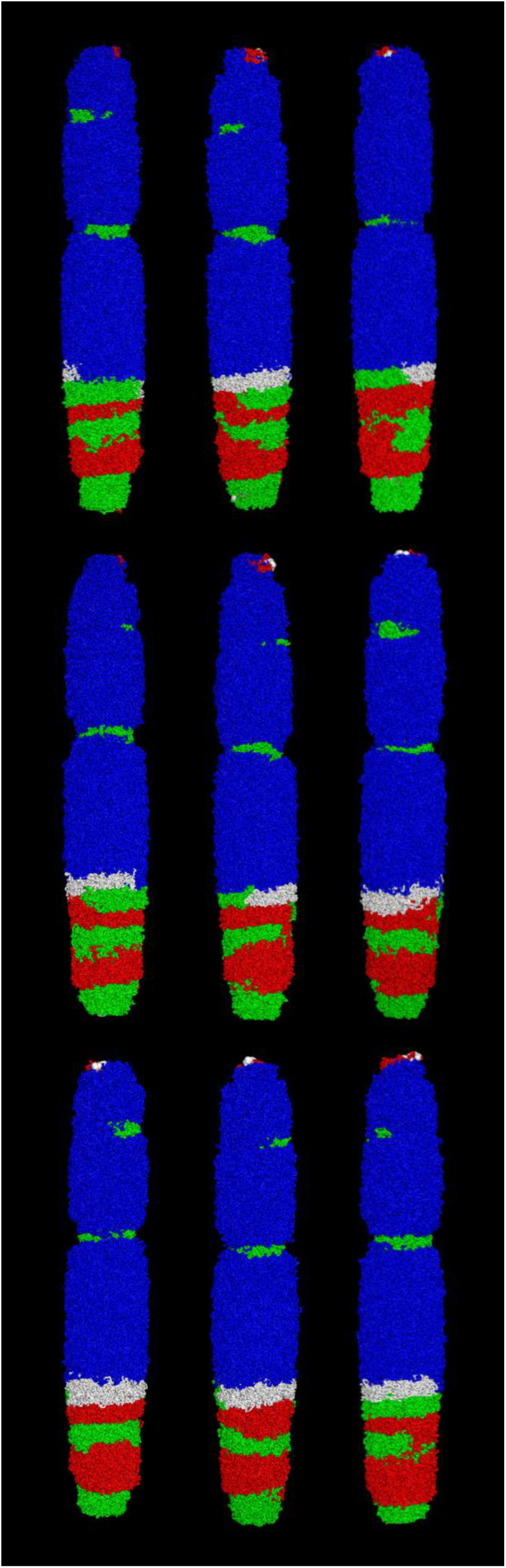
Nine replicates of the simulated Hi-C data-based polymer chromosome model. The simulations were based on the turn lengths calculated from the experimental Hi-C data. The different patterns arose from the varying loop distributions. Their positions and sizes were chosen at random according to an exponential distribution to reach the same average loop size.

**Figure S15.**
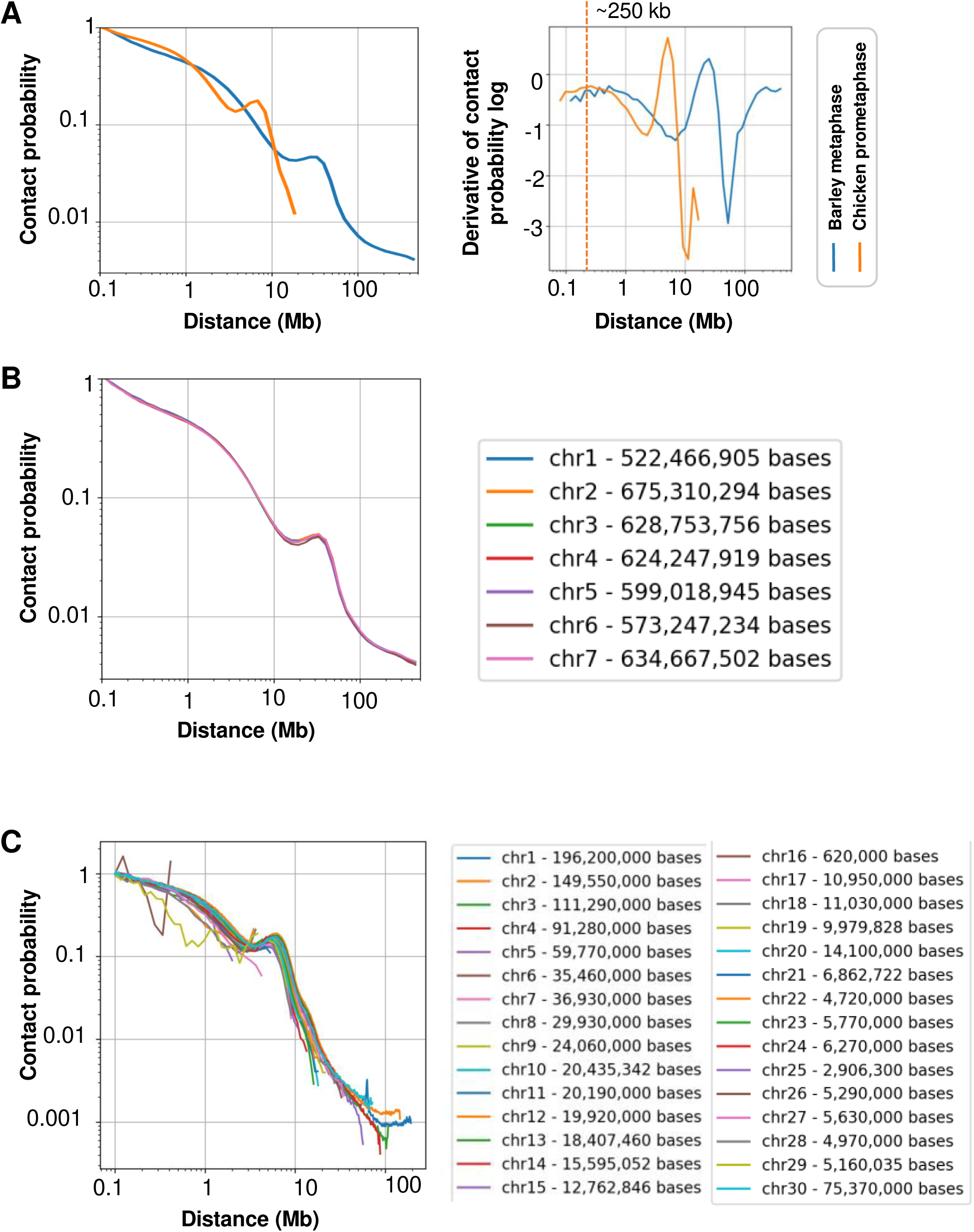
Comparison of Hi-C-based contact probabilities of mitotic chromosomes in chicken vs. barley. **(A)** Contact probabilities of barley chromosome 5H at metaphase and chicken chromosome 1 at prometaphase (Gibcus et al. 2018). The right diagram shows the derivative of the log of the same contact probabilities, clearly indicating the bump position. The dashed line marks the average size of the nested loops **(B)** Contact probabilities of all barley chromosomes at metaphase and **(C)** all chicken chromosomes at prometaphase (Gibcus et al. 2018). All probabilities were normalized to be equal to 1 at 100 kb.

**Figure S16:**
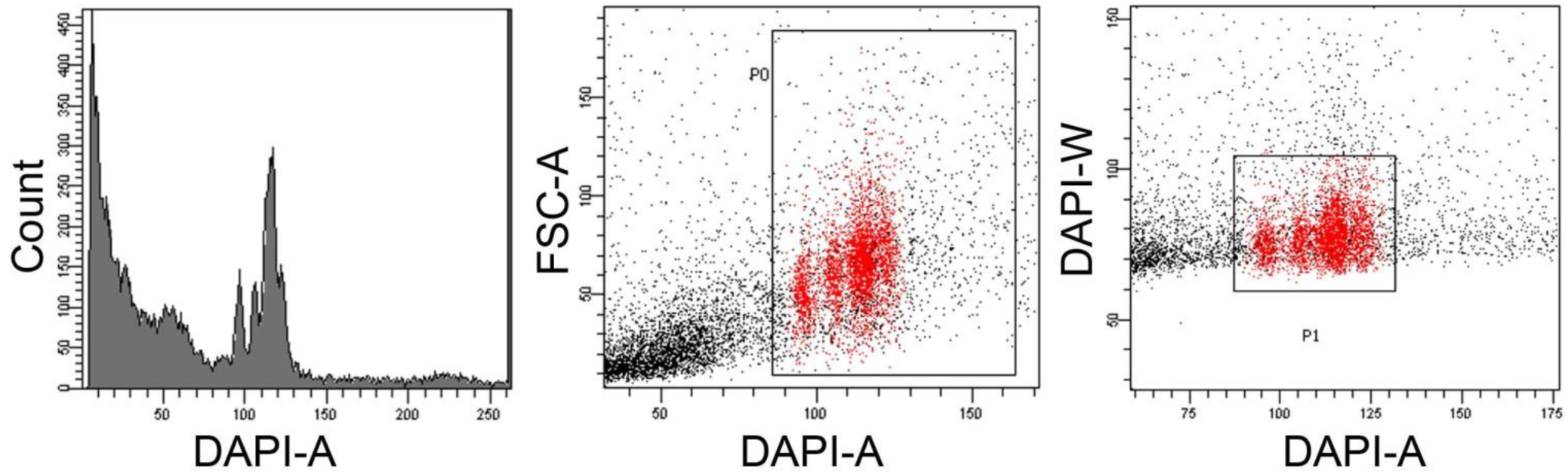
Flow sorting of barley metaphase chromosomes. Flow karyotype of barley cv. Morex. Initial gating was set on relative DAPI fluorescence vs. forward scatter parameters (P0); the dependent sorting gate was drawn based on the DAPI-area vs. DAPI-width scatter plot (P1).

**Table S1:**
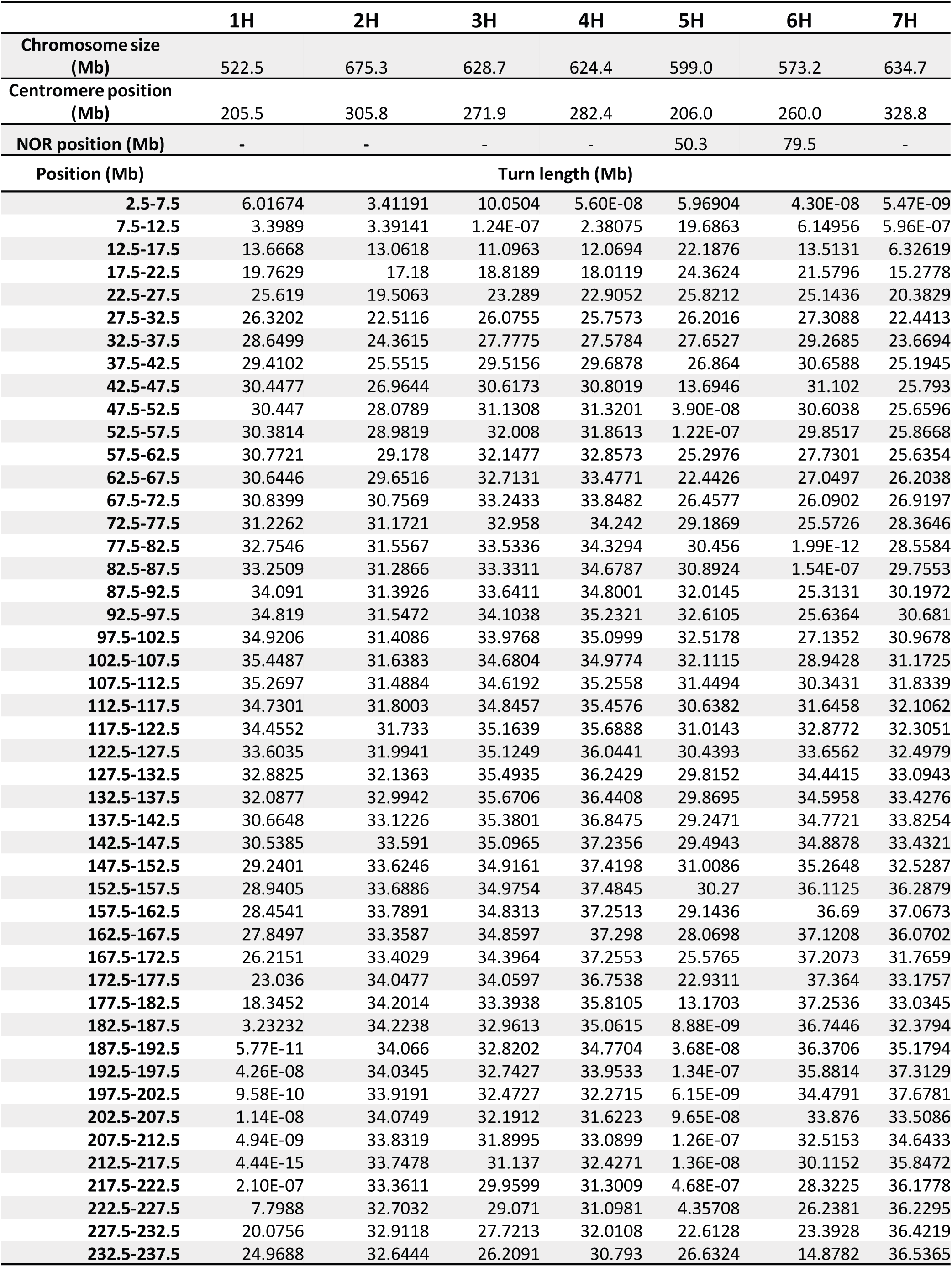
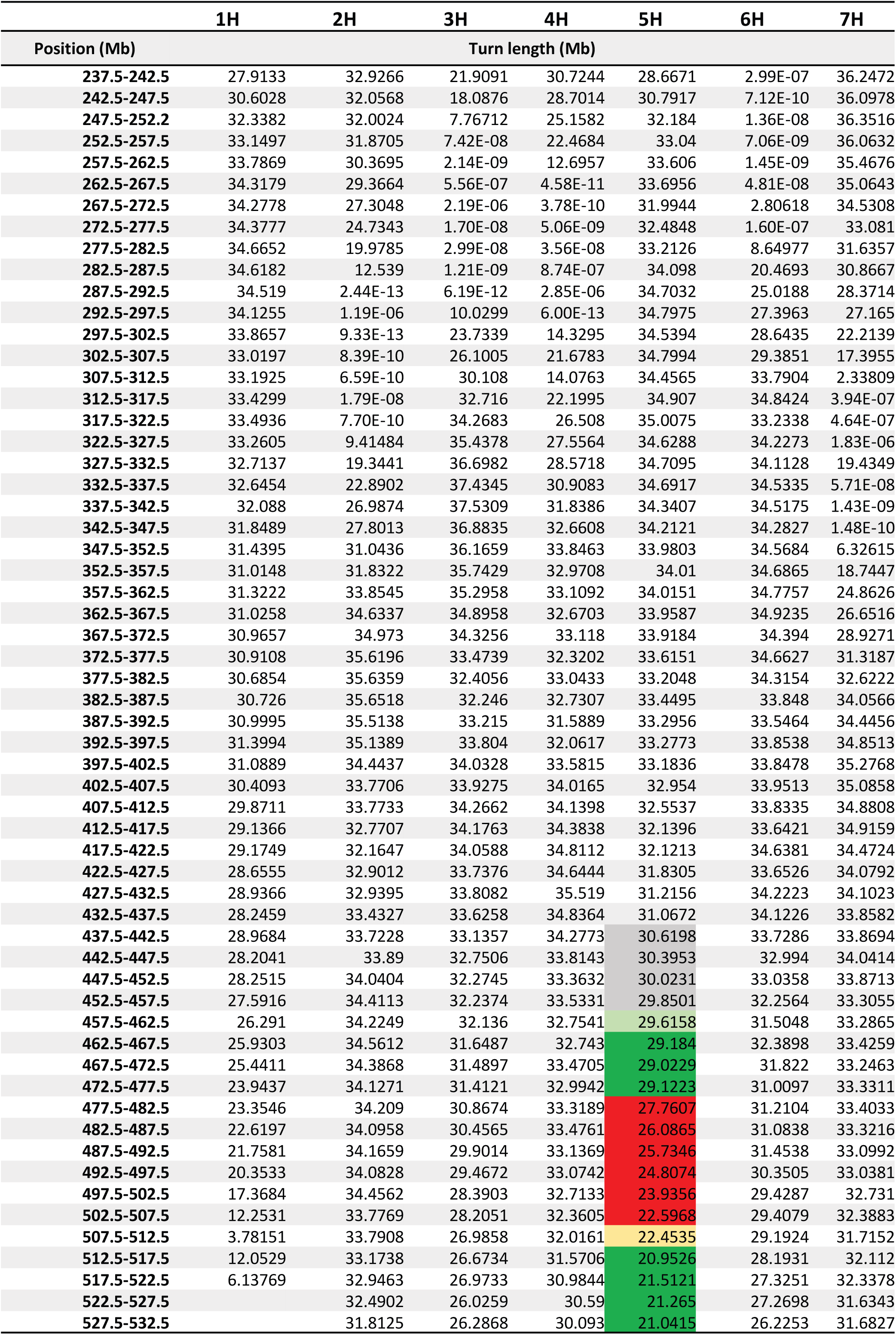
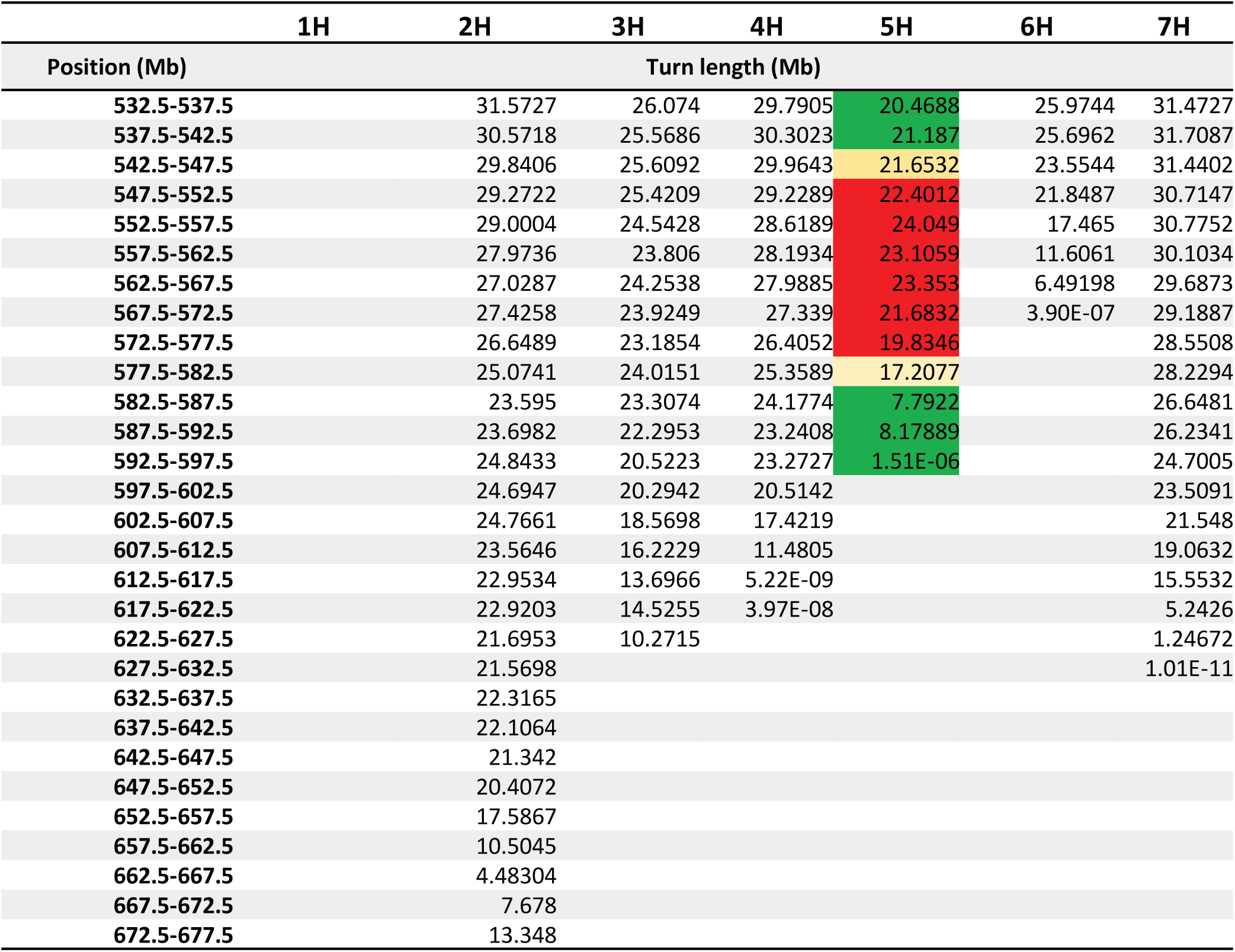
Values of local turn length of each chromosome calculated for 5-Mb bins, as described in the supplementary materials and Figure S4. These values are graphically shown in Figure 1E. The first three lines display the sequence length of each chromosome and the positions of the primary (centromere) and secondary constriction (NOR) at the short arms of 5H and 6H, as predicted from Hi-C data.

|  | 1H | 2H | 3H | 4H | 5H | 6H | 7H |
| --- | --- | --- | --- | --- | --- | --- | --- |
| Chromosome size (Mb) | 522.5 | 675.3 | 628.7 | 624.4 | 599.0 | 573.2 | 634.7 |
| Centromere position (Mb) | 205.5 | 305.8 | 271.9 | 282.4 | 206.0 | 260.0 | 328.8 |
| NOR position (Mb) | - | - | - | - | 50.3 | 79.5 | - |
| Position (Mb) | Turn length (Mb) |  |  |  |  |  |  |
| 2.5-7.5 | 6.01674 | 3.41191 | 10.0504 | 5.60E-08 | 5.96904 | 4.30E-08 | 5.47E-09 |
| 7.5-12.5 | 3.3989 | 3.39141 | 1.24E-07 | 2.38075 | 19.6863 | 6.14956 | 5.96E-07 |
| 12.5-17.5 | 13.6668 | 13.0618 | 11.0963 | 12.0694 | 22.1876 | 13.5131 | 6.32619 |
| 17.5-22.5 | 19.7629 | 17.18 | 18.8189 | 18.0119 | 24.3624 | 21.5796 | 15.2778 |
| 22.5-27.5 | 25.619 | 19.5063 | 23.289 | 22.9052 | 25.8212 | 25.1436 | 20.3829 |
| 27.5-32.5 | 26.3202 | 22.5116 | 26.0755 | 25.7573 | 26.2016 | 27.3088 | 22.4413 |
| 32.5-37.5 | 28.6499 | 24.3615 | 27.7775 | 27.5784 | 27.6527 | 29.2685 | 23.6694 |
| 37.5-42.5 | 29.4102 | 25.5515 | 29.5156 | 29.6878 | 26.864 | 30.6588 | 25.1945 |
| 42.5-47.5 | 30.4477 | 26.9644 | 30.6173 | 30.8019 | 13.6946 | 31.102 | 25.793 |
| 47.5-52.5 | 30.447 | 28.0789 | 31.1308 | 31.3201 | 3.90E-08 | 30.6038 | 25.6596 |
| 52.5-57.5 | 30.3814 | 28.9819 | 32.008 | 31.8613 | 1.22E-07 | 29.8517 | 25.8668 |
| 57.5-62.5 | 30.7721 | 29.178 | 32.1477 | 32.8573 | 25.2976 | 27.7301 | 25.6354 |
| 62.5-67.5 | 30.6446 | 29.6516 | 32.7131 | 33.4771 | 22.4426 | 27.0497 | 26.2038 |
| 67.5-72.5 | 30.8399 | 30.7569 | 33.2433 | 33.8482 | 26.4577 | 26.0902 | 26.9197 |
| 72.5-77.5 | 31.2262 | 31.1721 | 32.958 | 34.242 | 29.1869 | 25.5726 | 28.3646 |
| 77.5-82.5 | 32.7546 | 31.5567 | 33.5336 | 34.3294 | 30.456 | 1.99E-12 | 28.5584 |
| 82.5-87.5 | 33.2509 | 31.2866 | 33.3311 | 34.6787 | 30.8924 | 1.54E-07 | 29.7553 |
| 87.5-92.5 | 34.091 | 31.3926 | 33.6411 | 34.8001 | 32.0145 | 25.3131 | 30.1972 |
| 92.5-97.5 | 34.819 | 31.5472 | 34.1038 | 35.2321 | 32.6105 | 25.6364 | 30.681 |
| 97.5-102.5 | 34.9206 | 31.4086 | 33.9768 | 35.0999 | 32.5178 | 27.1352 | 30.9678 |
| 102.5-107.5 | 35.4487 | 31.6383 | 34.6804 | 34.9774 | 32.1115 | 28.9428 | 31.1725 |
| 107.5-112.5 | 35.2697 | 31.4884 | 34.6192 | 35.2558 | 31.4494 | 30.3431 | 31.8339 |
| 112.5-117.5 | 34.7301 | 31.8003 | 34.8457 | 35.4576 | 30.6382 | 31.6458 | 32.1062 |
| 117.5-122.5 | 34.4552 | 31.733 | 35.1639 | 35.6888 | 31.0143 | 32.8772 | 32.3051 |
| 122.5-127.5 | 33.6035 | 31.9941 | 35.1249 | 36.0441 | 30.4393 | 33.6562 | 32.4979 |
| 127.5-132.5 | 32.8825 | 32.1363 | 35.4935 | 36.2429 | 29.8152 | 34.4415 | 33.0943 |
| 132.5-137.5 | 32.0877 | 32.9942 | 35.6706 | 36.4408 | 29.8695 | 34.5958 | 33.4276 |
| 137.5-142.5 | 30.6648 | 33.1226 | 35.3801 | 36.8475 | 29.2471 | 34.7721 | 33.8254 |
| 142.5-147.5 | 30.5385 | 33.591 | 35.0965 | 37.2356 | 29.4943 | 34.8878 | 33.4321 |
| 147.5-152.5 | 29.2401 | 33.6246 | 34.9161 | 37.4198 | 31.0086 | 35.2648 | 32.5287 |
| 152.5-157.5 | 28.9405 | 33.6886 | 34.9754 | 37.4845 | 30.27 | 36.1125 | 36.2879 |
| 157.5-162.5 | 28.4541 | 33.7891 | 34.8313 | 37.2513 | 29.1436 | 36.69 | 37.0673 |
| 162.5-167.5 | 27.8497 | 33.3587 | 34.8597 | 37.298 | 28.0698 | 37.1208 | 36.0702 |
| 167.5-172.5 | 26.2151 | 33.4029 | 34.3964 | 37.2553 | 25.5765 | 37.2073 | 31.7659 |
| 172.5-177.5 | 23.036 | 34.0477 | 34.0597 | 36.7538 | 22.9311 | 37.364 | 33.1757 |
| 177.5-182.5 | 18.3452 | 34.2014 | 33.3938 | 35.8105 | 13.1703 | 37.2536 | 33.0345 |
| 182.5-187.5 | 3.23232 | 34.2238 | 32.9613 | 35.0615 | 8.88E-09 | 36.7446 | 32.3794 |
| 187.5-192.5 | 5.77E-11 | 34.066 | 32.8202 | 34.7704 | 3.68E-08 | 36.3706 | 35.1794 |
| 192.5-197.5 | 4.26E-08 | 34.0345 | 32.7427 | 33.9533 | 1.34E-07 | 35.8814 | 37.3129 |
| 197.5-202.5 | 9.58E-10 | 33.9191 | 32.4727 | 32.2715 | 6.15E-09 | 34.4791 | 37.6781 |
| 202.5-207.5 | 1.14E-08 | 34.0749 | 32.1912 | 31.6223 | 9.65E-08 | 33.876 | 33.5086 |
| 207.5-212.5 | 4.94E-09 | 33.8319 | 31.8995 | 33.0899 | 1.26E-07 | 32.5153 | 34.6433 |
| 212.5-217.5 | 4.44E-15 | 33.7478 | 31.137 | 32.4271 | 1.36E-08 | 30.1152 | 35.8472 |
| 217.5-222.5 | 2.10E-07 | 33.3611 | 29.9599 | 31.3009 | 4.68E-07 | 28.3225 | 36.1778 |
| 222.5-227.5 | 7.7988 | 32.7032 | 29.071 | 31.0981 | 4.35708 | 26.2381 | 36.2295 |
| 227.5-232.5 | 20.0756 | 32.9118 | 27.7213 | 32.0108 | 22.6128 | 23.3928 | 36.4219 |
| 232.5-237.5 | 24.9688 | 32.6444 | 26.2091 | 30.793 | 26.6324 | 14.8782 | 36.5365 |
| Position (Mb) | Turn length (Mb) |  |  |  |  |  |  |
| 237.5-242.5 | 27.9133 | 32.9266 | 21.9091 | 30.7244 | 28.6671 | 2.99E-07 | 36.2472 |
| 242.5-247.5 | 30.6028 | 32.0568 | 18.0876 | 28.7014 | 30.7917 | 7.12E-10 | 36.0978 |
| 247.5-252.2 | 32.3382 | 32.0024 | 7.76712 | 25.1582 | 32.184 | 1.36E-08 | 36.3516 |
| 252.5-257.5 | 33.1497 | 31.8705 | 7.42E-08 | 22.4684 | 33.04 | 7.06E-09 | 36.0632 |
| 257.5-262.5 | 33.7869 | 30.3695 | 2.14E-09 | 12.6957 | 33.606 | 1.45E-09 | 35.4676 |
| 262.5-267.5 | 34.3179 | 29.3664 | 5.56E-07 | 4.58E-11 | 33.6956 | 4.81E-08 | 35.0643 |
| 267.5-272.5 | 34.2778 | 27.3048 | 2.19E-06 | 3.78E-10 | 31.9944 | 2.80618 | 34.5308 |
| 272.5-277.5 | 34.3777 | 24.7343 | 1.70E-08 | 5.06E-09 | 32.4848 | 1.60E-07 | 33.081 |
| 277.5-282.5 | 34.6652 | 19.9785 | 2.99E-08 | 3.56E-08 | 33.2126 | 8.64977 | 31.6357 |
| 282.5-287.5 | 34.6182 | 12.539 | 1.21E-09 | 8.74E-07 | 34.098 | 20.4693 | 30.8667 |
| 287.5-292.5 | 34.519 | 2.44E-13 | 6.19E-12 | 2.85E-06 | 34.7032 | 25.0188 | 28.3714 |
| 292.5-297.5 | 34.1255 | 1.19E-06 | 10.0299 | 6.00E-13 | 34.7975 | 27.3963 | 27.165 |
| 297.5-302.5 | 33.8657 | 9.33E-13 | 23.7339 | 14.3295 | 34.5394 | 28.6435 | 22.2139 |
| 302.5-307.5 | 33.0197 | 8.39E-10 | 26.1005 | 21.6783 | 34.7994 | 29.3851 | 17.3955 |
| 307.5-312.5 | 33.1925 | 6.59E-10 | 30.108 | 14.0763 | 34.4565 | 33.7904 | 2.33809 |
| 312.5-317.5 | 33.4299 | 1.79E-08 | 32.716 | 22.1995 | 34.907 | 34.8424 | 3.94E-07 |
| 317.5-322.5 | 33.4936 | 7.70E-10 | 34.2683 | 26.508 | 35.0075 | 33.2338 | 4.64E-07 |
| 322.5-327.5 | 33.2605 | 9.41484 | 35.4378 | 27.5564 | 34.6288 | 34.2273 | 1.83E-06 |
| 327.5-332.5 | 32.7137 | 19.3441 | 36.6982 | 28.5718 | 34.7095 | 34.1128 | 19.4349 |
| 332.5-337.5 | 32.6454 | 22.8902 | 37.4345 | 30.9083 | 34.6917 | 34.5335 | 5.71E-08 |
| 337.5-342.5 | 32.088 | 26.9874 | 37.5309 | 31.8386 | 34.3407 | 34.5175 | 1.43E-09 |
| 342.5-347.5 | 31.8489 | 27.8013 | 36.8835 | 32.6608 | 34.2121 | 34.2827 | 1.48E-10 |
| 347.5-352.5 | 31.4395 | 31.0436 | 36.1659 | 33.8463 | 33.9803 | 34.5684 | 6.32615 |
| 352.5-357.5 | 31.0148 | 31.8322 | 35.7429 | 32.9708 | 34.01 | 34.6865 | 18.7447 |
| 357.5-362.5 | 31.3222 | 33.8545 | 35.2958 | 33.1092 | 34.0151 | 34.7757 | 24.8626 |
| 362.5-367.5 | 31.0258 | 34.6337 | 34.8958 | 32.6703 | 33.9587 | 34.9235 | 26.6516 |
| 367.5-372.5 | 30.9657 | 34.973 | 34.3256 | 33.118 | 33.9184 | 34.394 | 28.9271 |
| 372.5-377.5 | 30.9108 | 35.6196 | 33.4739 | 32.3202 | 33.6151 | 34.6627 | 31.3187 |
| 377.5-382.5 | 30.6854 | 35.6359 | 32.4056 | 33.0433 | 33.2048 | 34.3154 | 32.6222 |
| 382.5-387.5 | 30.726 | 35.6518 | 32.246 | 32.7307 | 33.4495 | 33.848 | 34.0566 |
| 387.5-392.5 | 30.9995 | 35.5138 | 33.215 | 31.5889 | 33.2956 | 33.5464 | 34.4456 |
| 392.5-397.5 | 31.3994 | 35.1389 | 33.804 | 32.0617 | 33.2773 | 33.8538 | 34.8513 |
| 397.5-402.5 | 31.0889 | 34.4437 | 34.0328 | 33.5815 | 33.1836 | 33.8478 | 35.2768 |
| 402.5-407.5 | 30.4093 | 33.7706 | 33.9275 | 34.0165 | 32.954 | 33.9513 | 35.0858 |
| 407.5-412.5 | 29.8711 | 33.7733 | 34.2662 | 34.1398 | 32.5537 | 33.8335 | 34.8808 |
| 412.5-417.5 | 29.1366 | 32.7707 | 34.1763 | 34.3838 | 32.1396 | 33.6421 | 34.9159 |
| 417.5-422.5 | 29.1749 | 32.1647 | 34.0588 | 34.8112 | 32.1213 | 34.6381 | 34.4724 |
| 422.5-427.5 | 28.6555 | 32.9012 | 33.7376 | 34.6444 | 31.8305 | 33.6526 | 34.0792 |
| 427.5-432.5 | 28.9366 | 32.9395 | 33.8082 | 35.519 | 31.2156 | 34.2223 | 34.1023 |
| 432.5-437.5 | 28.2459 | 33.4327 | 33.6258 | 34.8364 | 31.0672 | 34.1226 | 33.8582 |
| 437.5-442.5 | 28.9684 | 33.7228 | 33.1357 | 34.2773 | 30.6198 | 33.7286 | 33.8694 |
| 442.5-447.5 | 28.2041 | 33.89 | 32.7506 | 33.8143 | 30.3953 | 32.994 | 34.0414 |
| 447.5-452.5 | 28.2515 | 34.0404 | 32.2745 | 33.3632 | 30.0231 | 33.0358 | 33.8713 |
| 452.5-457.5 | 27.5916 | 34.4113 | 32.2374 | 33.5331 | 29.8501 | 32.2564 | 33.3055 |
| 457.5-462.5 | 26.291 | 34.2249 | 32.136 | 32.7541 | 29.6158 | 31.5048 | 33.2865 |
| 462.5-467.5 | 25.9303 | 34.5612 | 31.6487 | 32.743 | 29.184 | 32.3898 | 33.4259 |
| 467.5-472.5 | 25.4411 | 34.3868 | 31.4897 | 33.4705 | 29.0229 | 31.822 | 33.2463 |
| 472.5-477.5 | 23.9437 | 34.1271 | 31.4121 | 32.9942 | 29.1223 | 31.0097 | 33.3311 |
| 477.5-482.5 | 23.3546 | 34.209 | 30.8674 | 33.3189 | 27.7607 | 31.2104 | 33.4033 |
| 482.5-487.5 | 22.6197 | 34.0958 | 30.4565 | 33.4761 | 26.0865 | 31.0838 | 33.3216 |
| 487.5-492.5 | 21.7581 | 34.1659 | 29.9014 | 33.1369 | 25.7346 | 31.4538 | 33.0992 |
| 492.5-497.5 | 20.3533 | 34.0828 | 29.4672 | 33.0742 | 24.8074 | 30.3505 | 33.0381 |
| 497.5-502.5 | 17.3684 | 34.4562 | 28.3903 | 32.7133 | 23.9356 | 29.4287 | 32.731 |
| 502.5-507.5 | 12.2531 | 33.7769 | 28.2051 | 32.3605 | 22.5968 | 29.4079 | 32.3883 |
| 507.5-512.5 | 3.78151 | 33.7908 | 26.9858 | 32.0161 | 22.4535 | 29.1924 | 31.7152 |
| 512.5-517.5 | 12.0529 | 33.1738 | 26.6734 | 31.5706 | 20.9526 | 28.1931 | 32.112 |
| 517.5-522.5 | 6.13769 | 32.9463 | 26.9733 | 30.9844 | 21.5121 | 27.3251 | 32.3378 |
| 522.5-527.5 |  | 32.4902 | 26.0259 | 30.59 | 21.265 | 27.2698 | 31.6343 |
| 527.5-532.5 |  | 31.8125 | 26.2868 | 30.093 | 21.0415 | 26.2253 | 31.6827 |
| Position (Mb) | Turn length (Mb) |  |  |  |  |  |  |
| 532.5-537.5 |  | 31.5727 | 26.074 | 29.7905 | 20.4688 | 25.9744 | 31.4727 |
| 537.5-542.5 |  | 30.5718 | 25.5686 | 30.3023 | 21.187 | 25.6962 | 31.7087 |
| 542.5-547.5 |  | 29.8406 | 25.6092 | 29.9643 | 21.6532 | 23.5544 | 31.4402 |
| 547.5-552.5 |  | 29.2722 | 25.4209 | 29.2289 | 22.4012 | 21.8487 | 30.7147 |
| 552.5-557.5 |  | 29.0004 | 24.5428 | 28.6189 | 24.049 | 17.465 | 30.7752 |
| 557.5-562.5 |  | 27.9736 | 23.806 | 28.1934 | 23.1059 | 11.6061 | 30.1034 |
| 562.5-567.5 |  | 27.0287 | 24.2538 | 27.9885 | 23.353 | 6.49198 | 29.6873 |
| 567.5-572.5 |  | 27.4258 | 23.9249 | 27.339 | 21.6832 | 3.90E-07 | 29.1887 |
| 572.5-577.5 |  | 26.6489 | 23.1854 | 26.4052 | 19.8346 |  | 28.5508 |
| 577.5-582.5 |  | 25.0741 | 24.0151 | 25.3589 | 17.2077 |  | 28.2294 |
| 582.5-587.5 |  | 23.595 | 23.3074 | 24.1774 | 7.7922 |  | 26.6481 |
| 587.5-592.5 |  | 23.6982 | 22.2953 | 23.2408 | 8.17889 |  | 26.2341 |
| 592.5-597.5 |  | 24.8433 | 20.5223 | 23.2727 | 1.51E-06 |  | 24.7005 |
| 597.5-602.5 |  | 24.6947 | 20.2942 | 20.5142 |  |  | 23.5091 |
| 602.5-607.5 |  | 24.7661 | 18.5698 | 17.4219 |  |  | 21.548 |
| 607.5-612.5 |  | 23.5646 | 16.2229 | 11.4805 |  |  | 19.0632 |
| 612.5-617.5 |  | 22.9534 | 13.6966 | 5.22E-09 |  |  | 15.5532 |
| 617.5-622.5 |  | 22.9203 | 14.5255 | 3.97E-08 |  |  | 5.2426 |
| 622.5-627.5 |  | 21.6953 | 10.2715 |  |  |  | 1.24672 |
| 627.5-632.5 |  | 21.5698 |  |  |  |  | 1.01E-11 |
| 632.5-637.5 |  | 22.3165 |  |  |  |  |  |
| 637.5-642.5 |  | 22.1064 |  |  |  |  |  |
| 642.5-647.5 |  | 21.342 |  |  |  |  |  |
| 647.5-652.5 |  | 20.4072 |  |  |  |  |  |
| 652.5-657.5 |  | 17.5867 |  |  |  |  |  |
| 657.5-662.5 |  | 10.5045 |  |  |  |  |  |
| 662.5-667.5 |  | 4.48304 |  |  |  |  |  |
| 667.5-672.5 |  | 7.678 |  |  |  |  |  |
| 672.5-677.5 |  | 13.348 |  |  |  |  |  |

**Table S2:** Coordinates of the oligo-FISH probes (named after different birds) adjacent to the 5HL subtelomere based on the barley cv. Morex v2 genome assembly.

| Probe | No. oligos | Start (Mb) | End (Mb) | Length (Mb) | Average Turn size (Mb) | Number of turns* |
| --- | --- | --- | --- | --- | --- | --- |
| Stork | 33987 | 442.1 | 459.9 | 17.7 | 30.1±0.4 | 0.6 |
| Eagle | 35062 | 459.9 | 477.5 | 17.6 | 29.2±0.2 | 0.6 |
| Ostrich | 91061 | 477.4 | 511.8 | 34.4 | 24.8±1.8 | 1.4 |
| Rhea | 105984 | 511.8 | 545.3 | 33.5 | 21.3±0.5 | 1.58 |
| Moa | 108831 | 545.3 | 580.2 | 34.9 | 21.6±2.1 | 1.62 |
| Flamingo | 62181 | 580.1 | 599.0 | 18.8 | 8.3±6.1 | 2.3 |

\* Length of the probe divided by the average turn size in its region.

**Table S3:** Volumes of barley satellite chromosomes 5H and 6H (identified based on their secondary constriction and centromere positions) and NOR-free (1H-4H, 7H) chromosomes. The measurements were done on polyacrylamide embedded spatially preserved chromosomes.

| Chromosome | n | Mean±SD (µm<sup>3</sup>) |
| --- | --- | --- |
| 5H | 2 | 49.5±2.3 |
| 6H | 3 | 55.0±12.8 |
| 1H-4H, 7H | 10 | 56.1±9.8 |
| 1H-7H | 15 | 55.0±10.1 |

**Table S4:** Statistics of the metaphase Hi-C datasets. Seven libraries were analyzed: three replicates from interphase libraries, three replicates from metaphase libraries, and one metaphase library generated with an alternative protocol, which yielded fewer unspecific contacts, for comparison with polymer modeling. For each library, the total number of contact reads (all reads), percentage of PCR duplicated reads (duplicated), number of non-duplicated reads with a mapping quality (MAPQ) of at least 10 (high-quality reads), number of intra-chromosome contacts separated by more than 1 kb (1 kb separation), number of intra-chromosome contacts separated by more than 1 Mb (1 Mb separation), and number of inter-chromosome contacts (inter-chromosome) were computed. * percentage of all reads: ** percentage of reads relative to the number of high-quality reads; *** percentage of reads separated by more than 1 kb.

| LIBRARY | ALL<br>READS | DUPLICATED<br>(%*) | HIGH-QUALITY<br>READS<br>(%*) | 1KB<br>SEPARATION<br>(%**) | 1MB<br>SEPARATION<br>(%***) | INTER-CHROMOSOME<br>(%***) |
| --- | --- | --- | --- | --- | --- | --- |
| INTERPHASE_1 | 260,968,878 | 13.21 | 39,558,947<br>(15.16) | 16,036,382<br>(40.54) | 7,110,811<br>(44.34) | 4,138,138<br>(25.8) |
| INTERPHASE_2 | 204,856,338 | 1.19 | 31,097,110<br>(15.18) | 18,557,444<br>(59.68) | 8,604,498<br>(46.37) | 4,567,769<br>(24.61) |
| INTERPHASE_3 | 219,294,615 | 1.49 | 36,292,924<br>(16.55) | 35,634,600<br>(98.19) | 12,389,518<br>(34.77) | 18,915,266<br>(53.08) |
| METAPHASE_1 | 509,196,403 | 6.06 | 91,401,051<br>(17.95) | 62,357,229<br>(68.22) | 12,764,657<br>(20.47) | 48,635,089<br>(77.99) |
| METAPHASE_2 | 735,344,585 | 8.28 | 92,941,157<br>(12.64) | 69,241,898<br>(74.5) | 23,853,444<br>(34.45) | 41,427,153<br>(59.83) |
| METAPHASE_3 | 579,559,917 | 7.09 | 80,301,755<br>(13.86) | 60,884,103<br>(75.82) | 17,350,884<br>(28.5) | 40,922,614<br>(67.21) |
| METAPHASE_AGAR | 122,636,722 | 50.74 | 12,194,950<br>(9.94) | 9,809,014<br>(80.43) | 4,771,893<br>(48.65) | 3,254,155<br>(33.17) |

## Movies

**Movie 1.** Bump position along chromosome 5H, indicating the turn sizes. Placement of the bump calculated for 119 different positions in the sequence of chromosome 5H separated by 5 Mb.

**Movie 2.** The 14 somatic barley metaphase chromosomes labeled by FISH, as described in Figure 2B-D. The movie clearly shows that the oligo-FISH labeled 5HL region is also labeled inside of the chromosome arms.

**Movie 3.** Slices of a chromosome 5H SIM image stack shown in ortho view (Figure 6B), indicating the complete FISH labeling of the inner chromatin.

**Movie 4:** Spatial minor chromatin fiber intermingling of adjacent chromonema turns at chromosome arm 5HL labeled in different colors by oligo-FISH (Figure S10).

**Movie 5.** Spatial surface rendering of chromosome 5H (Figure 6C) showing that FISH signals are present within the whole chromosome volume.

**Movie 6.** Spatial arrangements of telomeres and subtelomeres (Figure 6E).

**Movie 7.** Spatial arrangement of the oligo-FISH and EdU labeled region of the left 5H chromosome (*) shown in Figure 4.

**Movie 8.** Spatial arrangement of the oligo-FISH and EdU labeled region of the right 5H chromosome (**) shown in Figure 4.

**Movie 9.** Single SIM image stack slices of a DAPI-labeled 6H chromosome showing a network of minor ∼80 nm looped chromatin fibers free of large cavities throughout the chromosome arms. Within the centromere and NOR, straight ∼80 nm chromatin fibers occur.

**Movie 10.** Spatial organization of the same 6H chromosome shown in Movie 9.

**Movie 11.** Animation of the polymer simulation model shown in Figure 6D for a single FISH-labeled 5H chromatid.

**Movie 12.** Polymer simulation of two helical chromonema half-turns (red and green) of a metaphase chromatid within a cylinder.

## Notes

### Competing Interest Statement

The authors have declared no competing interest.

### Summary of Updates

Text revised. Supplemental files added.

